# Two-stage Linked Component Analysis for Joint Decomposition of Multiple Biologically Related Data Sets

**DOI:** 10.1101/2021.03.22.435728

**Authors:** Huan Chen, Brian Caffo, Genevieve Stein-O’Brien, Jinrui Liu, Ben Langmead, Carlo Colantuoni, Luo Xiao

## Abstract

Integrative analysis of multiple data sets has the potential of fully leveraging the vast amount of high throughput biological data being generated. In particular such analysis will be powerful in making inference from publicly available collections of genetic, transcriptomic and epigenetic data sets which are designed to study shared biological processes, but which vary in their target measurements, biological variation, unwanted noise, and batch variation. Thus, methods that enable the joint analysis of multiple data sets are needed to gain insights into shared biological processes that would otherwise be hidden by unwanted intra-data set variation. Here, we propose a method called two-stage linked component analysis (2s-LCA) to jointly decompose multiple biologically related experimental data sets with biological and technological relationships that can be structured into the decomposition. The consistency of the proposed method is established and its empirical performance is evaluated via simulation studies. We apply 2s-LCA to jointly analyze four data sets focused on human brain development and identify meaningful patterns of gene expression in human neurogenesis that have shared structure across these data sets.

## 1. Introduction

### 1.1 Background

Heterogeneous data of various types are now being collected from scientific experiments and/or large scale health initiatives. Thus, it is of scientific interest to harness the collective discovery potential of such complex and growing data resources. For example, in biomedical research, high-dimensional gene expression and epigenetic data are produced to gain insight into cellular processes and disease mechanisms (Stein-O’Brien *and others*, 2019). Another example is in the study of Alzheimer’s disease where recent efforts have been focusing on combining brain imaging data, genetic data, as well as clinical outcomes in predicting disease (Nathoo *and others*, 2019). A third example is a large study profiling different states of wellness, where genetic, proteomic and metabolic data among other types of data are collected (Gao *and others*, 2020). Together, the different types of data provide a more comprehensive picture which has the potential of better characterizing and optimizing what it is to be healthy. In addition to these examples of studies with multimodal data collection, many public repositories are full of experimental data from single modalities that are related biologically. Therefore, in recent years, there has been growing interests as well as demands for understanding and utilizing these data in an integrative way (Lock *and others*, 2013; Yang and Michailidis, 2016; Li and Jung, 2017; Gaynanova and Li, 2019; Gao *and others*, 2020).

### 1.2 Motivating Data

Advances in RNA sequencing technologies have produced a large amount of data creating unprecedented opportunities for scientific discovery through data integration. We put together four experimental paradigms used to study brain development: i. van de Leemput *and others* (2014); *ii*. Yao *and others* (2017); iii. BrainSpan (2011); and *iv*. Nowakowski *and others* (2017). Specifically, we use two *in vitro* data sets from cultured human pluripotent stem cells subjected to neural differentiation, and two *in vivo* data sets of brain tissue across varying ages of human fetal development. Within the *in vitro* and *in vivo* studies two different RNA sequencing technologies were used: the two bulk studies extracted RNA from all the cells in tissue samples and expression levels were measured in aggregate for each tissue sample, while in the two single-cell studies, the RNA from each cell in a tissue sample was extracted and quantified individually. All four data sets focus on the creation of neurons from neural precursor cells across these times and hence are highly interrelated. However, there are sample specific details that introduce experiment specific variation beyond the expected technical batch effects. In the van de Leemput *and others* (2014) bulk data set, data were collected at 9 time points across 77 days of neural differentiation with a total number of 24 *in vitro* samples. The Yao *and others* (2017) data set was collected to model the production of human cortical neurons *in vitro*. There are about 2.7 thousand cells collected over across 54 days during *in vitro*. In the BrainSpan (2011) bulk data set, there are a total of 35 *in vivo* brain tissue samples. The Nowakowski *and others* (2017) data set contains around 3.5 thousand single cells across different ages in the fetal brain. For visualization of these data sets, see the NeMO Analytics portal at https://nemoanalytics.org/p?l=ChenEtAl2021&g=DCX, where the gEAR platform is leveraged to construct an integrated gene expression data viewer including all data sets used in the report (Orvis *and others*, 2021).

The primary goal of assembling these different data sets together is to define molecular dynamics that are common to all of them and are central in the creation of human neocortical neurons. In addition, we are interested in learning what cellular processes during *in vivo* development of the brain can be recapitulated in the *in vitro* systems and what processes are only present in the *in vivo* data. To achieve this goal, we need a new joint model that can properly model the two-way design exhibited by the four data sets and separate the different cellular processes.

### 1.3 Existing Literature

We distinguish two distinct types of data structure for multiple data sets: multi-view data and linked data. For multi-view data, e.g., multi-omics data, different sets of features are measured for each subject or unit with subject being the linking entity across data sets. For linked data, e.g., gene expression data for different types of cancer patients in The Cancer Genome Atlas (TCGA), the same set of features are measured for different groups of subjects. When data sets are organized such that each column corresponds to one sample, the former can be called vertically linked data as the data sets can be aligned vertically, while the latter can be called horizontally linked data as the data sets can be aligned horizontally (Richardson *and others*, 2016). For both types of data structures, a central goal of statistical analysis is to identify meaningful decomposition of variations across data sets.

Canonical correlation analysis (CCA) is a useful method for extracting common variation across multi-view data. Various variants of CCA for high dimensional data sets have been proposed; see, e.g., Gao *and others* (2020) and Min and Long (2020). However, CCA ignores potentially substantial variation present only in individual data sets. To remedy this, the joint and individual variations explained (JIVE) method (Lock *and others*, 2013) and subsequent variants of JIVE, e.g., AJIVE (Feng *and others*, 2018), were developed to identify common variation shared across multiple data sets as well as individual variation specific to each data set. Going further, the structural learning and integrative decomposition (SLIDE) method (Gaynanova and Li, 2019) also allows variations that are shared by only a subset of data sets. Other relevant works include Li and Jung (2017), Li *and others* (2018) and Argelaguet *and others* (2018).

A pioneering method for linked data is common principal component analysis (Flury, 1984, PCA), which can identify the same set of orthogonal eigenvectors shared across data sets. CPCA was extended to extract shared signal subspace present across data sets (Flury, 1987) and to determine the number of shared components (Wang *and others*, 2020). Another extension based on matrix decomposition, population value decomposition (Crainiceanu *and others*, 2011) extends CPCA to matrix-valued data, e.g., neuroimaging and electrophysiology data.

We mention a few other related works. Lock *and others* (2020) and Park and Lock (2020) jointly analyzed multiple data sets for heterogeneous groups of objects with heterogeneous feature sets. Kallus *and others* (2019) developed a data adaptive method which allows structure in data to be shared across an arbitrary subset of views and cohorts. Gao *and others* (2020) and Wang and Allen (2021) considered clustering problems for multi-view data. Li *and others* (2019) proposed a regression model with multi-view data as covariates.

### 1.4 Our Contribution

We propose a joint decomposition model for linked multiple data sets with a general design, e.g., the 2 by 2 factorial design in our motivating data, thus extending the aforementioned CPCA method which assumes the linked multiple data sets have a one-way design. The goal of the joint decomposition is to identify signal subspaces that are either shared or not shared across data sets. Similar in spirit to the underlying model for the SLIDE method for multi-view data, our model allows for common signal subspace that are shared by all data sets, partially shared signal subspaces that are shared by only subsets of all data sets, and individual signal subspaces specific to each data set. It is important to note that, for our method, it is fixed signal subspaces rather than latent random scores that are shared across data sets. Another significant difference is that the existence of partially shared signal subspaces for the proposed model is determined by design, which is scientifically meaningful for our motivating data. While not designed explicitly for linked data and the use case described, other methods can also be used in this setting. The BIDIFAC+ method (Lock *and others*, 2020), an extension of the BIDIFAC method for bi-dimensional data (Park and Lock, 2020), could also accommodate partially shared subspaces by design for linked data. These methods were proposed for decomposing bi-dimensional data. Specifically, the BIDIFAC+ method decomposes data into the sum of matrices with varying structures and imposes matrix nuclear norm penalties for model selection. Additionally, the SLIDE method, while designed for multi-view data, may still be applied to linked data via transposing the data. Indeed, existing multi-view methods (e.g., JIVE) that are based on matrix decomposition can also be applicable for linked data. However, because SLIDE is a fully unsupervised method and due to the specific penalty it employs to learn different types of components, it is unclear how to directly incorporate design of components into SLIDE. Nevertheless, it is worth mentioning that SLIDE, BIDIFAC+, and the proposed model allow partially shared components.

Our model estimation consists of a simple and intuitive two-stage procedure and is built on existing literature in principal component analysis. From a computational perspective, it is straightforward and avoids complex and computationally challenging optimization methods, e.g., large matrix decomposition and iterative optimization methods as in BIDIFAC+ and SLIDE. In particular, the proposed method avoids joint estimation of dimensions of different types of spaces, which can be challenging for both BIDFIFAC+ and SLIDE as is shown in the simulation study. We also study the theoretic properties of the proposed method for a general class of decomposition models. Specifically, we establish the consistency of the proposed method for estimating the signal subspaces. In particular, we prove the consistency of the proposed method for estimating the dimensions of the signal subspaces. Note that most existing methods in the literature lack theoretic justification for the choice of dimensions (or ranks of matrices).

The rest of the paper is organized as follows. In Section 2, we first describe our model for linked data sets with a 2 by 2 design, as motivated by our data example, and then extend the model for data sets with a general design. A two-step model estimation method is given in the Section. In Section 3, we study the theoretic properties of the proposed model estimation method. In Section 4, we carry out a simulation study to evaluate the empirical performance of our proposed method and compare with existing methods. In Section 5, we apply the proposed method to the motivating data and also carry out a validation study using additional data sets. Finally, in Section 6, we discuss how our work can be extended, e.g., for high-dimensional data.

## 2. Models

### 2.1 Model Formulation

First, consider a 2 by 2 factorial design, as in our motivating data, but later generalize the model for more generic designs. Let *i* be the index for the cells’ system, with *i* = 1 for *in vivo* and *i* = 2 for *in vitro*. Similarly, let *j* be the index for the RNA sequencing technology, with *j* = 1 for single cell and *j* = 2 for bulk. Let 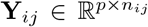 be the (*i*, *j*)th data set consisting of *n_ij_* tissue samples or cells of dimension *p*. simplicity, hereafter we shall use sample to refer to either tissue sample or cell. Denote by 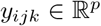 the *k*th sample for the (*i*, *j*)th data set so that **Y**_*ij*_ = [*y*_*ij*i_,…, *y_ijn_ij__*]. Thus, the samples are of the same dimension for all data sets and they are assumed independent between the data sets and from each other.

Consider a two-way latent factor model,

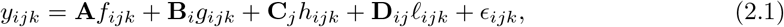

where 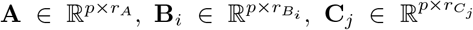 and 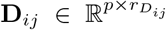 are fixed orthonormal matrices of components with associated random scores 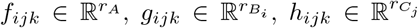 and 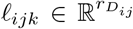, respectively. The vector 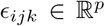 consists of uncorrelated, mean zero error terms, with variance 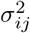. We assume that all random scores have zero means and cov(*f_ijk_*) = **Σ**_*fij*_, cov(*g_ijk_*) = **Σ**_*gij*_, cov(*h_ijk_*) = **Σ**_h_ij__, cov(*ℓ_ijk_*) = **Σ**_*ℓij*_, with positive diagonal elements in all covariances. In particular, we assume without loss of generality that **Σ**_*f*11_, **Σ**_*gi*1_ (*i* = 1, 2), **Σ**_*h*1*j*_ (*j* = 1, 2), and **Σ**_*ℓij*_ (*i* = 1, 2, *j* = 1, 2) are diagonal matrices so that the matrices can be identified. The other covariance matrices can be non-diagonal, implying that components do not have to align component-wisely across the data sets, but the space spanned by them are the same. Finally, the random terms {*f_ijk_*, *g_ijk_*, *h_ijk_*, *ℓ_ijk_*, *ε_ijk_*} are assumed uncorrelated across *i*, *j*, and *k*.

For model identifiability, we impose the condition that the orthogonal matrices **A**, **B**_1_, **B**_2_, **C**_1_, **C**_2_, **D**_*j*_ (1 ⩽ *i*, *j* ⩽ 2) are also mutually orthogonal. This condition ensures that the components are not movable, e.g., components in **A** can not be transferred to other matrices without changing the model.

Model (2.1) can be interpreted as a multivariate ANOVA model. To elaborate, we use the notation col(·) to denote the column space of a matrix. Then, col(**A**) is the space of common components, the space of variation that is shared by all data sets. Next, col(**B**_1_) is the space of partially shared components specific to *in vivo* system, i.e., the two *in vivo* data sets. Thus, it is orthogonal to both the common components, i.e., col(**A**), and the space of variation specific to the *in vitro* system, i.e., col(**B**_2_). Third, col(**C**_1_) is the space of partially shared components specific to the single cell sequencing technology and col(**C**_2_) is for the bulk sequencing method. Finally, col(**D**_*ij*_) are the individual components specific to each data set.

For data set (*i*, *j*), model (2.1) is a latent factor model with the signal space being the column space of **U**_*ij*_ = [**A**, **B**_*i*_, **C**_*j*_, **D**_*ij*_], which can be denoted by col(**U**_*ij*_).

### 2.2 Model Estimation

We propose a two-stage estimation method, called 2s-LCA. The first stage is to estimate the signal space from each data set separately using principal component analysis (PCA). The second stage combines the signal spaces to extract the common subspace, the partially shared subspaces and the individual subspaces, also using PCA. Note that the first stage of 2s-LCA is essentially the same as the first step of AJIVE (Feng *and others*, 2018), which first extracts signal spaces from each data set and then decomposes the variations of the obtained latent scores into common variation and individual variation.

For the first stage, the key is to estimate the number of components in each data set, i.e., the dimension of col(Ujo). We use the BEMA method (Ke *and others*, 2021) (see supplementary material for detail), which fits a distribution to the eigenvalues (putatively) arising from noise using the observed bulk eigenvalues and then determines the number of eigenvalues arising from signals via extrapolation of the fitted distribution. This method yields consistent estimation and worked well in our simulations. Denote by **Û**_*ij*_ the matrices of the estimated eigenvectors associated with the top eigenvalues from the sample covariance of data set (*i*, *j*).

For the second stage, we extract the common subspace, the partially shared subspaces, and the individual subspaces, sequentially. Assume that we extract the subspaces in the following order: **A** → **B**_1_ → **B**_2_ → **C**_1_ → **C**_2_ → **D**_11_ → **D**_12_ → **D**_21_ → **D**_22_. Here, to simplify notation, we use the matrices to denote the subspaces.

The estimation of the common subspace is based on the following observation. Let 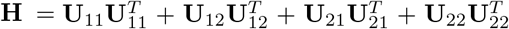. Then it is easy to show that for any eigenvector of **H**, a^*T*^**H**a equals 4 if a ∈ col(**A**) and is at most 2 otherwise. Thus, there exists a natural gap of eigenvalues associated with the common subspace and those of other subspaces for **H**. From a theoretical perspective, any cut-off value between 2 and 4 for the eigenvalues may lead to consistent estimation. In practice, a scree plot of the eigenvalues of 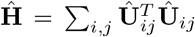 can be used to determine the dimension of the common subspace, i.e. considering the relative amount of variation explained among the top eigenvalues. Once the dimension of the common subspace is determined, the eigenvectors of **Ĥ** associated with these top eigenvalues give us an estimate of the space of common components. Let **Â** be the matrix with columns corresponding to the retained eigenvectors of **Ĥ**. Note that **Â** may not be a good estimate of **A** elementwise, rather, the notation is used to represent that col(**Â0**) is an estimate of col(**A**) and **Â** is one of many matrix representations of the former.

Next we estimate the partially shared subspaces. We first project **Û**_*ij*_ onto the orthogonal complement of **Â**, i.e., (**I** – **ÂÂ**^*T*^)**Û**_*ij*_. For simplicity, we still denote it by **Û**_ij_, but the common subspace has been removed. We now consider estimation of col(**B**_1_), the space of partially shared components corresponding to the *in vivo* system. With similar arguments as before, now the matrix 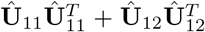 has one set of eigenvalues close to 2 and another set of eigenvalues close to or smaller than 1. Indeed, the theoretic counterpart of the above matrix has two sets of non-zero eigenvalues: one set of eigenvalues of 2 with associated eigenvectors forming the space col(**B**_1_) and another set of eigenvalues of 1 with associated eigenvectors from the individual space of either col(**D**_11_) or col(**D**_12_). Therefore, another scree plot of eigenvalues of 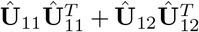 will determine the dimension of col(**B**_1_) and also give an estimate of the space.

Similar analysis can be used to estimate the other partially shared subspaces. For example, a spectral analysis of 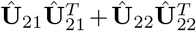 can be used to estimate col(**C**_1_). Further steps follow similarly and thus are omitted. To ensure the estimated spaces are orthogonal, orthogonal projection is always used after each estimation of the partially shared subspaces. After the projection of **Û**_*ij*_ onto the complement of estimates of col(**B**_*i*_) and col(**C**_*j*_), the remaining space is an estimate of the individual space col(**D**_*ij*_).

A remaining problem lies in the determination of the order which subspaces are estimated. First, it is clear that components shared by more data sets should be prioritized, and estimated first. The problem then lies in the order to follow to extract the partially shared components. While this is not a problem in our theoretical analysis, in real data analysis, different orders produce different estimates, because of the orthogonal condition imposed on the model and the orthogonal projection used in model estimation. A reasonable method considers the quality of the data sets (e.g., the sample size) and the spaces of shared components with higher perceived quality prioritized in the estimation order.

### 2.3 Model and Estimation for General Design

We extend the above model for a two-way factorial design to a model with a general design. To accommodate the more general setting, the notation here is slightly different from the previous sections. Assume that there are *I* data sets each containing *n*_i_ samples. Denote by 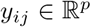 the *j*th sample for the *i*th data set. We consider a latent factor model with general design as follows:

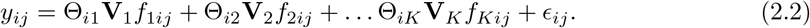

The orthonormal matrices 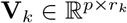 correspond to the *k*th space of components with associated scores, 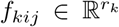. So the *k*th space is of dimension *r*_k_, which is unknown and needs to be determined. The indicator variables, Θ_*ik*_, denote if the *i*th data set contains the *k*th space of components, and 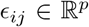 are random errors. Here the number of spaces, *K*, and the indicator variables are determined by design. For example, for the two-way factorial design considered above, *K* = 9. Two clear restrictions on the indicator variables are: (1) for each *k*, there exists at least one *i* such that Θ_*ik*_ = 1; (2) there does not exist *k* ≠ *k*′ with identical indicator values across all data sets. Let *η_k_* = ∑_i_ Θ_*ik*_, which is the number of data sets sharing the *k*th space of components. Without loss of generality, we assume that *η*_1_ ⩾… ⩾ *η_K_*.

We assume that all random scores have zero means and 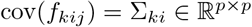 with positive diagonal elements in the covariance. In particular, for each *k*, there exists a pre-determined Θ*_ik_* = 1 such that Σ_*ki*_ is a diagonal matrix, i.e., the corresponding random vector *f_kij_* has uncorrelated elements. We also assume that all random terms are uncorrelated from each other and across *i* and *j*. For model identifiability, we impose that **V**_1_, **V**_2_,…, **V**_*K*_ are mutually orthogonal. Let 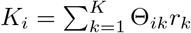, the total dimension of signal subspaces in data set *i*.

We now extend the model estimation proposed in the previous subsection. The first step remains the same and denote by **Û**_*i*_ the matrices of estimated eigenvectors. Consider the second step where the critical issue is the order of components to be extracted. The key idea is that spaces shared by more data sets should be extracted before those shared by fewer data sets. For each *k*, denote by 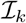 the index set for which if 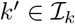, then Θ_*ik*′_ ⩽ Θ_*ik*_ and for at least one *i* the inequality is strict. Then let *α_k_* be the smallest element in 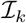. If 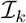 is empty then let *α_k_* be 0. Obviously, *α_k_* < *η_k_* for all *k*. Suppose we have estimated the first *k* spaces of shared components and have projected **Û**_*i*_ onto the orthogonal space of the previously extracted spaces. For simplicity, still denote those spaces by **Û**_*i*_. To estimate the (*k* + 1)th space, compute 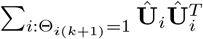 and its eigendecomposition. Let 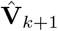 consists of the resulting eigenvectors with associated eigenvalues larger than (*η*_*k*+1_ + *α*_*k*+1_)/2. Then 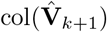 is our estimate of col(**V**_*k*+1_).

The above estimation procedure is unique if *η_k_* are all distinct. When there exists ties, then the order of estimation can be determined similarly as in the two-way design. One note is that our theoretical derivation remains valid no matter what order of estimation taken within the ties.

After obtaining the spaces of components, to further analyze or visualize the data, we project the data sets onto the estimated subspaces to obtain scores; see, e.g., our real data analysis.

## 3. Theoretical Properties

We establish the consistency of the proposed estimation methods for the general design proposed in (2). Specifically, we prove the consistency of 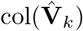 for estimating the signal space col(**V**_*k*_), for each *k*, which implies the consistency of dimension estimation for each signal subspace as well.

We use the following notation. For a square matrix **A**, denote by tr(**A**) the sum of diagonals in **A** and ||**A**||_2_ the operator norm of **A**. Let *r_e_*(**A**) = tr(**A**)/||**A**||_2_, *the effective rank* of **A**.

Consider the following assumptions.

### Assumption 3.1

*p*/*n* → *γ*, *γ* > 0 is a constant.

### Assumption 3.2

The minimal nonzero eigenvalue of **Σ**_*ki*_, denoted by *σ_ki_*, satisfies 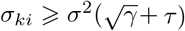, for a positive constant *τ* and *γ* in Assumption 3.1, for all *k* and *i*.

### Assumption 3.3

Assume that for each signal subspace **V**_*k*_, the random scores *f_kij_* ~ *N*(0, **Σ**_*ki*_), and the noise vector 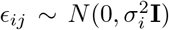, and that the random scores and noises are independent from each other and across subjects.

### Assumption 3.4

Let 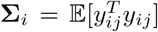 for *i* = 1,…, *n*. Assume that these covariance matrices have bounded effective ranks, i.e., *r_e_*(**Σ**_*i*_) ⩽ *C*, for all *i* and some fixed positive constant *C*.

Assumptions 3.1 and 3.2 are needed for the consistency of the BEMA method (Ke *and others*, 2021) for estimating the dimension of the signal space for each data matrix in the first step of model estimation. Assumption 3.2 means the signals can be separated from the noises. Assumptions 3.3 and 3.4 are needed to establish the consistency of the sample covariance matrix estimators (Bunea and Xiao, 2015). Assumption 3.4 means the signals have to be sufficiently strong so that they can be consistently estimated. When Assumption 3.4 is invalid, such as in the instance of a high dimensional spiked covariance matrix, additional sparsity assumptions are needed and then the sample covariances have to be replaced by sparsity-inducing estimators in our estimation method; see the discussion section for more details.

Let **V**, 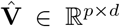 both have orthonormal columns, then the vector of *d* principal angles between their column spaces is given by (*cos*^-1^*σ*_1_,…, *cos*^-1^*σ_d_*)^*T*^, where *σ*_1_ ⩾… ⩾ *σ_d_* are the singular values of 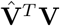. Let 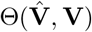 denote *d* × *d* diagonal matrix whose *j*th diagonal entry is the *j*th principal angle, and let 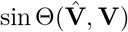 be defined entry-wise. Then, for two signal subspaces **V** and 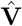, define convergence in space as 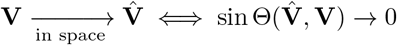.

Theorem 3.5 Under Model (2.2), if Assumptions 3.1 – 3.4 hold, then 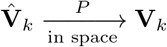, for all *k*.

The proof of Theorem 3.5 is provided in Section S.3 of the supplementary material. Note that this theorem also implies the consistency of the estimated dimension of each signal subspace.

## 4. Simulation Studies

### 4.1 Settings

Consider the two-way model (2.1) for which the spaces of common components, partially shared components, as well as individual components are all of dimension 2. We sample the orthonormal matrices (**A**, **B**_*i*_, **C**_*j*_, **D**_*jj*_) from a Stiefel manifold and generate the random scores and noises from normal distributions. For the diagonal covariance matrices (Σ_*f*11_, Σ_*g*11_, Σ_*h*11_, ι_*∑*11_, Σ_*ℓ*12_, ι_*∑*21_, Σ_*ℓ*22_), we sample the diagonals uniformly from the interval [1, 2] and then multiply them by the dimension *p*. To ensure desired signal to noise ratio as defined below, a scale parameter is multiplied to the diagonals. For the other covariance matrices, we randomly rotate Σ_*f*11_ to Σ_*fij*_ with a rotation angle *π*/3. Similarly, we randomly rotate Σ_*g*11_ and Σ_*h*11_ to Σ_*gij*_ and Σ_*hij*_ respectively with a rotation angle *π*/2. We generate noises *ε_ijk_* for each data set from *N*(0, *σ*^2^) with *σ*^2^ = 1.

We set the sample sizes *n_ij_* = *n* for all data sets and consider two cases. Case 1 is a high dimensional setting with *p* = 500, *n* = 500 and case 2 is a low dimensional setting with *p* = 50, *n* = 500. Finally, we define the signal to noise ratio (SNR) for the (*i*, *j*)th data set as SNR_*ij*_ = tr(cov(*y_ijk_*))/(*pσ*^2^) – 1. We set the same SNR for all data sets and use three different values of SNR, {0.2,1, 5}. Therefore, there are a total number of 6 model conditions. Under each model condition, we conduct 1000 simulations in a cluster computing environment.

We compare 2s-LCA with several existing methods. First, we fix the dimensions of all subspaces at their true values, i.e., 2, and compare 2s-LCA with JIVE (Lock *and others*, 2013), AJIVE (Feng *and others*, 2018), SLIDE (Gaynanova and Li, 2019), BIDIFAC (Park and Lock, 2020), and BIDIFAC+ (Lock *and others*, 2020) for subspace estimation. Then, we do the same as above except that the data are generated such that the variances of scores associated with individual subspaces are much larger than those for common and partially shared subspaces. Third and most importantly, we compare SLIDE, BIDIFAC+, and 2s-LCA without pre-fixing dimensions of the subspaces.

To evaluate the performance of methods, we use a metric called space alignment (SA), to measure the alignment of two spaces. For instance, if **Â** is the estimated common subspace and **A** is the population common subspace, then the SA between them is

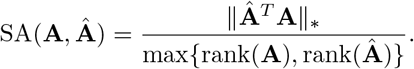

Here || · ||_*_ denotes the nuclear norm, which is invariant to matrix rotation. Note that SA always lies in [0,1], with 1 indicating that the two spaces are identical, while a 0 indicating that the two spaces are orthogonal to each other. We also compare the computational times of the proposed method and a few existing methods.

### 4.2 Simulation Results

We first summarize simulation results when the dimensions of the subspaces are fixed at their true values with further details provided in Section S.4.1 of the supplementary material. Because JIVE, AJIVE and BIDIFAC do not consider partially shared components, they do not perform well in this settings. On the contrary, SLIDE, BIDIFAC+, and 2s-LCA all perform well and have comparable values of SA across all model conditions.

However, when the variance of individual components are much larger than the common and partially shared components, both SLIDE and BIDIFAC+, can have difficulty recovering common and partially shared spaces. In contrast, 2s-LCA is capable of accurately estimating these spaces (Section S.4.2 of the supplementary material). A likely explanation is that both SLIDE and BIDIFAC+ focus on recovering the overall signal space, in this case dominated by the individual components, in the data while 2s-LCA prioritizes and first estimates common and partially shared components.

Now we compare SLIDE, BIDIFAC+, and 2s-LCA for model estimation without knowing the dimension of each space. As discussed in the Introduction, SLIDE does not allow direct specification of types of spaces and hence might be slightly disadvantaged compared with 2s-LCA and BIDIFAC+, as the latter two methods accommodate the specification of types of subspaces. Fig. 2 summarizes the performance of the methods under two model conditions. First, BIDIFAC+ tends to overestimate the dimensions of common and partially shared spaces, which negatively impacts SA. Second, SLIDE underestimates the dimensions of these spaces in low dimensional settings with small signal to noise ratios. Under both model conditions, 2s-LCA yields an accurate estimation of the dimensions of the subspaces, and results in high values for SA. Similar numerical results were found for the other four model conditions (Section S.4.3 of the supplementary materials). In particular, 2s-LCA seems capable of accurately estimating the dimensions of spaces correctly for most settings, except the high dimensional setting with a low SNR (Fig. S.16 of the supplementary material). However, it still has overall higher SA values than SLIDE and BIDIFAC+.

**Fig. 1:**
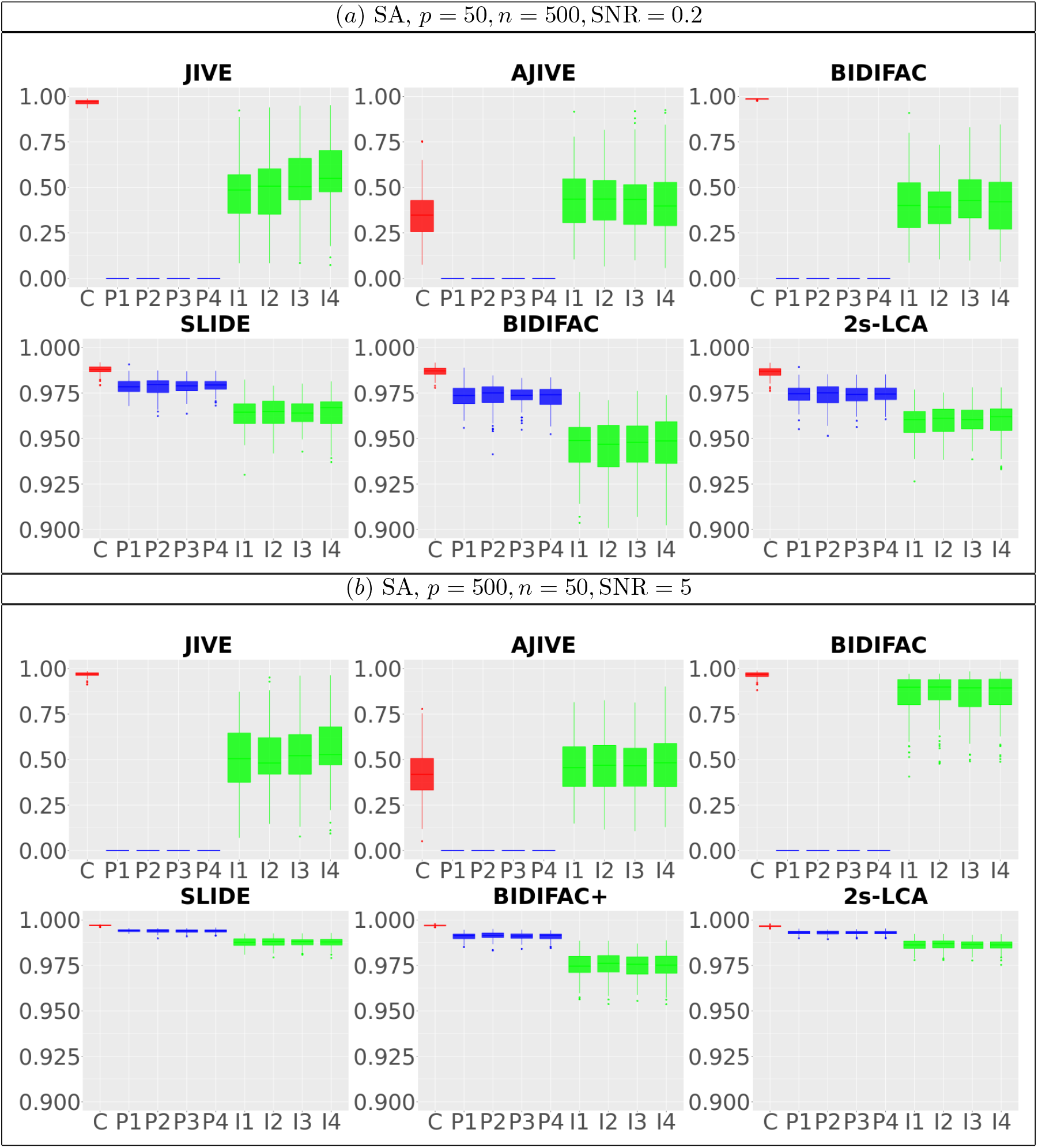
Comparisons between JIVE, AJIVE, BIDIFAC, SLIDE, BIDIFAC+ and 2s-LCA with known dimensions of spaces. (*a*) and (*b*) are both under the low dimensional setting with *p* = 50, *n* = 500, SNR = 0.2; (*c*) and (*d*) are under high dimensional setting with *p* = 500, *n* = 50, SNR = 5. For each setting, 1000 simulations are run.

**Fig. 2:**
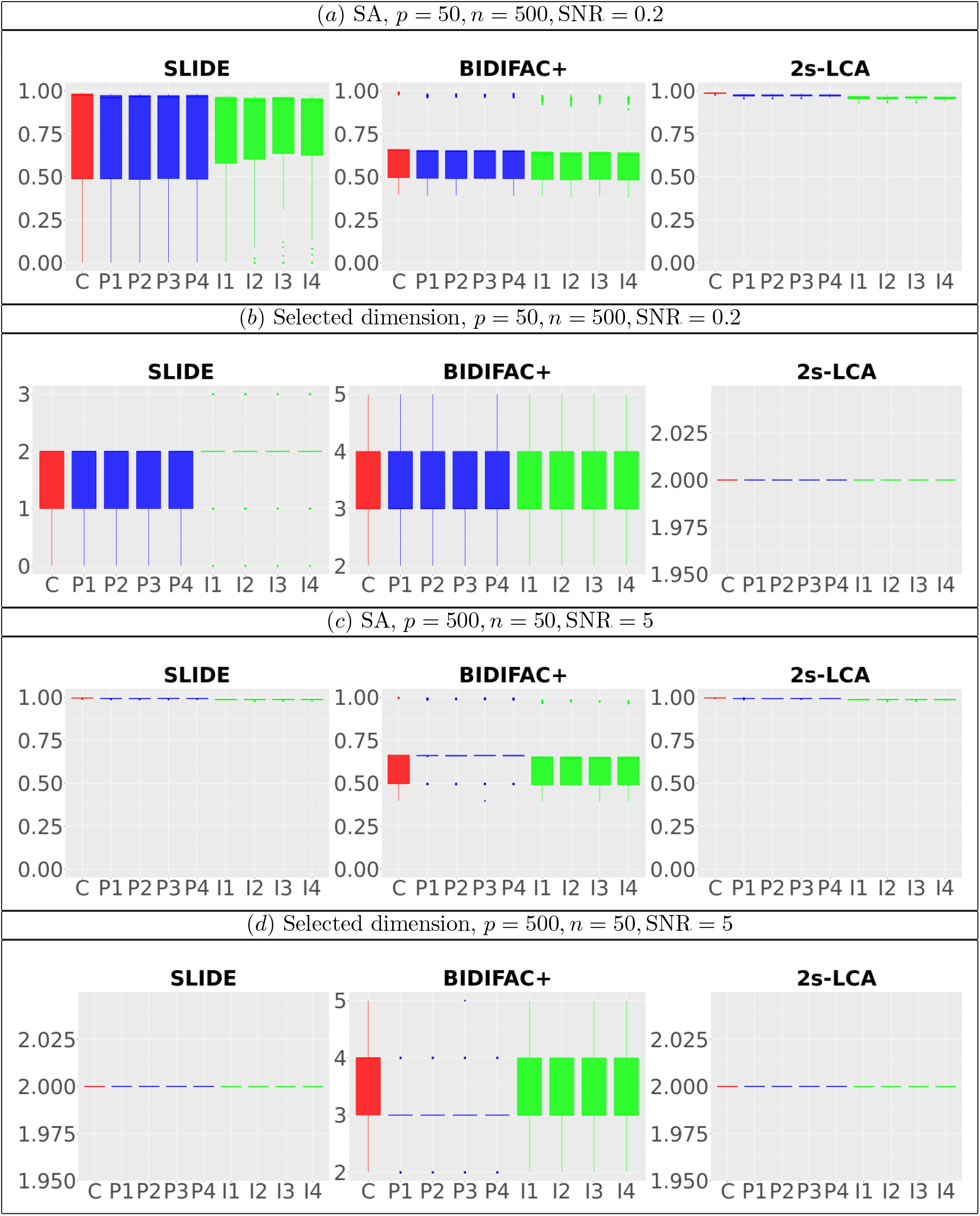
Comparisons between SLIDE, BIDIFAC+ and 2s-LCA. (*a*) and (*b*) are both under the low dimensional setting with *p* = 50, *n* = 500, SNR = 0.2; (*c*) and (*d*) are under high dimensional setting with *p* = 500, *n* = 50, SNR = 5. For each setting, 1000 simulations are run.

Lastly, we simulate the data as above except for setting the sizes of data as the same as that of the motivating data. We assume that the dimensions of spaces are known and fixed and run SLIDE, BIDIFAC+ and 2s-LCA. The proposed 2s-LCA takes about 32s (s.d. 16s) on average for one run while SLIDE takes about 6.5 hours (s.d. 3.0 hours) on average to reach convergence. We are unable to obtain convergent result for one single run after running the BIDIFAC+ code for 24 hours. We believe that the computational time of SLIDE can be potentially reduced by utilizing the inherent low rank property of the estimated signal matrix. As for BIDIFAC+, we believe its computational difficulties arise because it relies on eigendecompositions, which are difficult for high dimensional matrices. Consequently, BIDIFAC+ has a large RAM requirement, which may slow it down substantially if disk swapping becomes required. Finally, the iterative nature of both SLIDE and BIDIFAC+ may increase their computational time substantially.

## 5. Experimental Data Analysis

### 5.1 Analysis of Four Data Sets

We apply 2s-LCA to the four brain tissue data sets focused on brain development. As described in Section 1, the data sets were collected using two technologies, bulk and single cell RNA sequencing, and under two cellular environments, *in vivo* and *in vitro*. Thus, this exhibits a twoway design as in model (2.1), which includes one subspace of common components, four subspaces of partially shared components, and four subspaces of individual components. As two dimensional visualizations of the data are of interest, a natural choice of the number of components for each subspace is 2.

For 2s-LCA, normalization within each data set is needed to mitigate technical effects. For each data set, we center the expression value for each gene, center and scale expression level of each sample by its standard deviation across genes. For the second stage of 2s-LCA, we first estimate common components and then the partially shared components sequentially in the following order: *in vitro*, *in vivo*, single cell, and bulk.

Fig. 3 (*a*) shows the projection of every data set onto its top two eigenvectors–a separate analysis. We then apply 2s-LCA to jointly analyze the four data sets and obtain common, partially shared, and individual subspaces, each of which is of dimension 2. Then, we project every data set onto these subspaces to investigate biological processes associated with the scores; see Fig. 3 (*b*) for plots of the scores for common and partially shared subspaces. Note that the axes in each plot have different ranges and to facilitate interpretation, we have rotated the scores associated with the common subspace (top panel of Fig. 3 (*a*)) simultaneously for all four data sets.

**Fig. 3:**
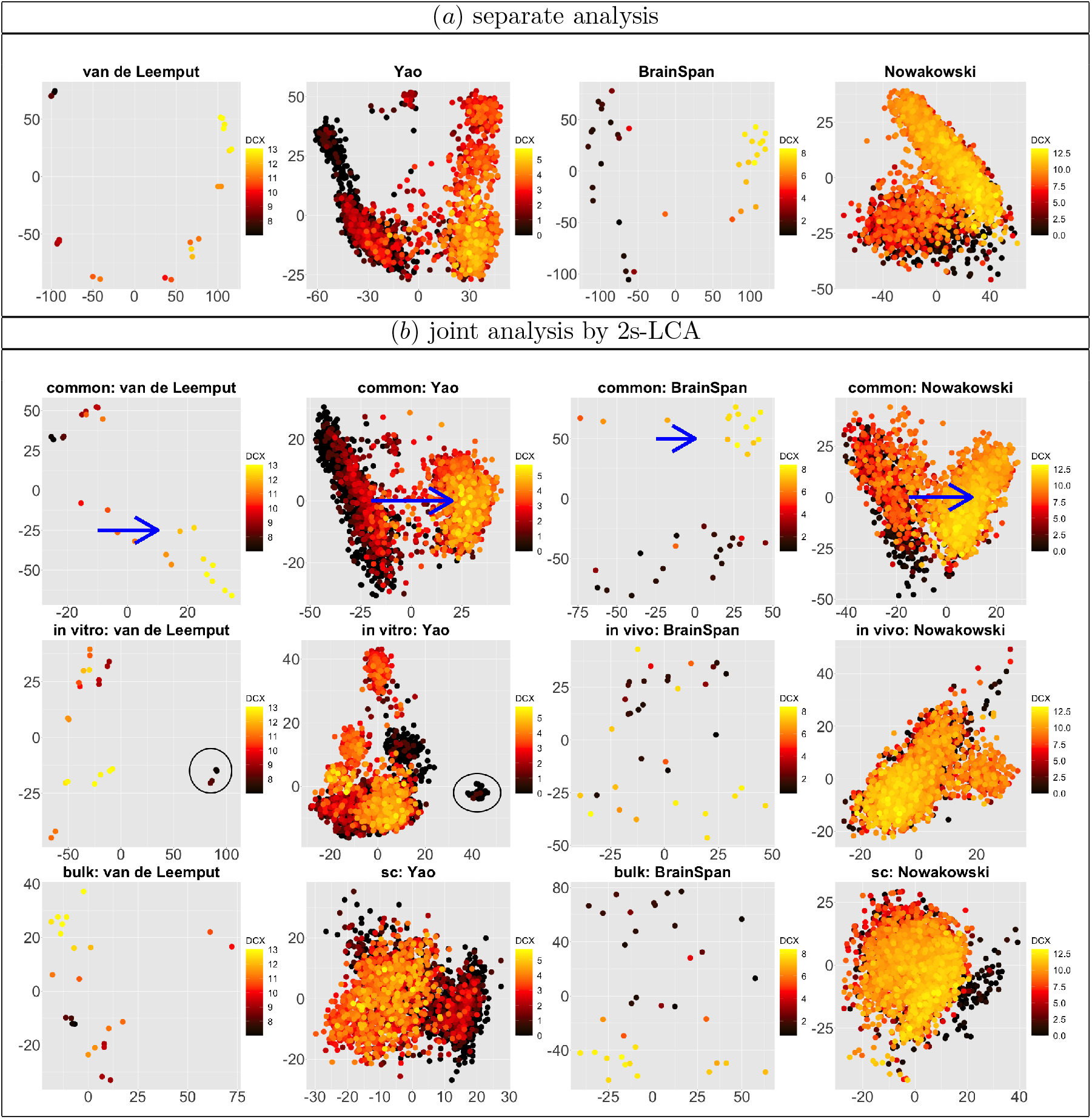
(*a*) Scatterplots of scores corresponding to the top two principal components of each data set by separate PCA and (*b*) Scatterplots of scores of each data set corresponding to common components (top panel), partially shared components associated with environments (middle panel: *in vitro* on the left two plots and *in vivo* on the right two plots), and partially shared components associated with technologies (bottom panel: bulk on the left two plots and single cell on the right two plots) by the proposed 2s-LCA. The 4 columns in both parts correspond to the data sets: (1) van de Leemput: *in vitro* + bulk; (2) Yao: *in vitro* + single cell; (3) BrainSpan: *in vivo* + bulk; and (4) Nowakowski: *in vivo* + single cell. Each point corresponds to either one tissue sample or one cell and is colored by the log_2_ transformed expression level of the DCX gene. Blue arrows indicate alignment of the first common component (x-axis in top row of panels in (b) with DCX expression (neurogenesis) in all four data sets. Black circles indicate pluripotent stem cells, which are present only in the *in vitro* data sets.

To evaluate and interpret results, we shall use the DCX gene, which is turned on as neural progenitors transition to being neurons. According to Liu (2011), “The DCX gene provides instructions for producing a protein called doublecortin. This protein is involved in the movement of nerve cells (neurons) to their proper locations in the developing brain, a process called neuronal migration.” The log_2_ transformed expression level of the DCX gene is used to color the cells, with the dark color indicating low expression level corresponding to neural progenitors and the yellow color indicating high expression level corresponding to neurons.

We first focus on the comparison of the separate components with the common components; see Fig. 3 (*a*) and the top row of panels in Fig. 3 (*b*). For the separate analysis, the plots show that cells with different expression levels of the DCX gene are clustered and separated; however, the extent to which the scores and gene loadings align across the data sets is unclear. In contrast to the separate analysis, 2s-LCA produces jointly derived components, allowing direct assessment of how effects across the different data sets align. The plots for common components show that cells with similar expression levels in the DCX gene tend to cluster together, which means that the local structure of the cells is preserved. In addition, by projecting the data sets onto the same subspace shared among them, a similar pattern can be observed among data sets, indicating shared global structure across the data sets. For example, the x-axis along which DCX expression increases during fetal development as neural progenitors become neurons (blue arrows) can be seen precisely aligned across the common components in each of the four data sets. Moreover, a visual comparison of the separate components with those of the proposed joint method suggests that some of the order defined in the separate components is preserved in the joint analysis (in the sense of the overall shape of the distribution of cells and clustering of cells with similar expression levels), consistent with the knowledge that these four diverse experiments capture common molecular elements of neurogenesis. In addition to the visual comparison, we also compute the ratio of the variances explained by the common components versus those explained by the separate analysis for each data set. The ratios are: (1) van de Leemput: 0.195; (2) Yao: 0.524; (3) BrainSpan: 0.382; and (4) Nowakowski: 0.538. The values indicate strong presence of common spaces shared by all data sets.

We have also evaluated the common components using the time of the samples: days of neural differentiation for the *in vitro* data sets and age (years) for the *in vivo* data sets; see Fig. 4 which is a recoloring of Fig. 3. Interestingly the *y*-axis (red arrows) appears to be a common temporal dimension orthogonal to the DCX expression component, which is along the *x*-axis in Fig. 3. As developmental time progresses, it appears that the cellular identities of the neural precursors and their post-mitotic neuronal progeny become less distinct, i.e. the clusters of low and high DCX expressing cells begin to merge. The cellular basis underlying this dimension is not known and might be of significant biological interest to explore further. In the top 50 gene specific loadings for the first common component, we identify, in addition to the DCX gene, many markers of nascent neurons, e.g., MYT1L and NRXN1, synaptic components, e.g., PSD2 and SYT1, and the channels SCN2A, SCN3A and SCN3B. Many of these identified genes have been implicated in human neurodevelopmental disorders (see supplementary table), which further demonstrates that this component parallels neurogensis in each of the four data sets.

**Fig. 4:**
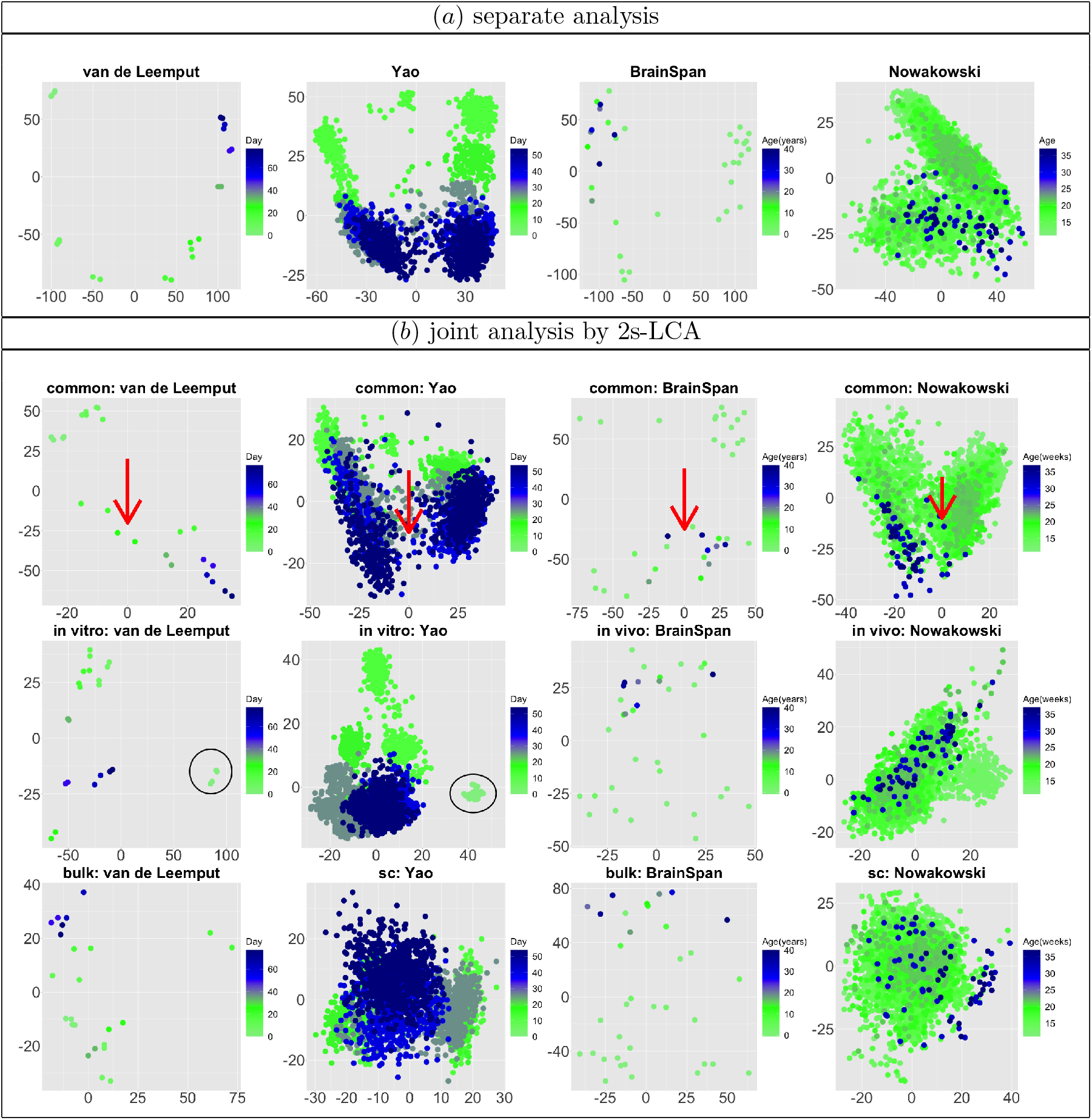
This figure is a re-coloring of the data shown in Fig. 3, in order to show effects across time. Each point corresponds to either one tissue sample or one cell and is colored by days of neural differentiation for the *in vitro* data sets and age in years or gestational weeks for the *in vivo* data sets. Red arrows indicate alignment of the second common component (*y*-axis in top row of panels in (*b*) with developmental time in all four data sets. Black circles indicate pluripotent stem cells, which are present only in the *in vitro* data sets.

We next evaluate the partially shared components and focus on the components shared only by the two *in vitro* data sets; see the left two plots on the second row of Fig. 3 (*b*). In both plots, those few cells circled in blue are pluripotent cells, which are present only in the *in vitro* data sets as they would have disappeared prior to the developmental time points measured in the *in vivo* system. This result demonstrates that the proposed joint decomposition method captures known biological effects unique to the *in vitro* experimental paradigm and not represented in the *in vivo* data sets. The top 10 gene specific loadings for the first *in vitro* component identifies pluripotent stem cells in the *in vitro* systems, as desired. Specifically, we find pluripotency markers: POU5F1, PRDM14, and CDH1, demonstrating the identification of a pluripotency transcriptional program common to the two *in vitro* data sets.

Finally, we also applied the BIDIFAC+ and SLIDE methods to the data sets. In a shared cluster computing environment, it took SLIDE about 1 day to yield convergent result. The results from SLIDE are are provided in the supplementary material. The results have similar apparent scientific validity as 2s-LCA. The results for BIDIFAC+ are unavailable, as we could not obtain convergent results in several days, again likely due to the requirements of the eigenvalue decomposition. We believe the computational efficiency of 2s-LCA is a key aspect to the model. For example, for this data convergence took only a two minutes.

### 5.2 A Validation Study

To validate the common components defined by the joint 2s-LCA decomposition (top row of panels in Fig. 3 (*b*) and Fig. 4 (*b*), we use 8 additional single cell RNA-seq data sets from 5 studies, see the supplementary material for details about these studies and the data sets. We project the additional data sets on the found common and *in vitro* components through the “projectR” package (Sharma *and others*, 2020). Projection of these data onto the common components clearly demonstrates recapitulation of the biological effect captured in the first component that is aligned with DCX expression and neurogenesis, and to a lesser extent the second that is aligned with time (Fig. 5 (*a*) and (*b*), blue and red arrows, respectively).

**Fig. 5:**
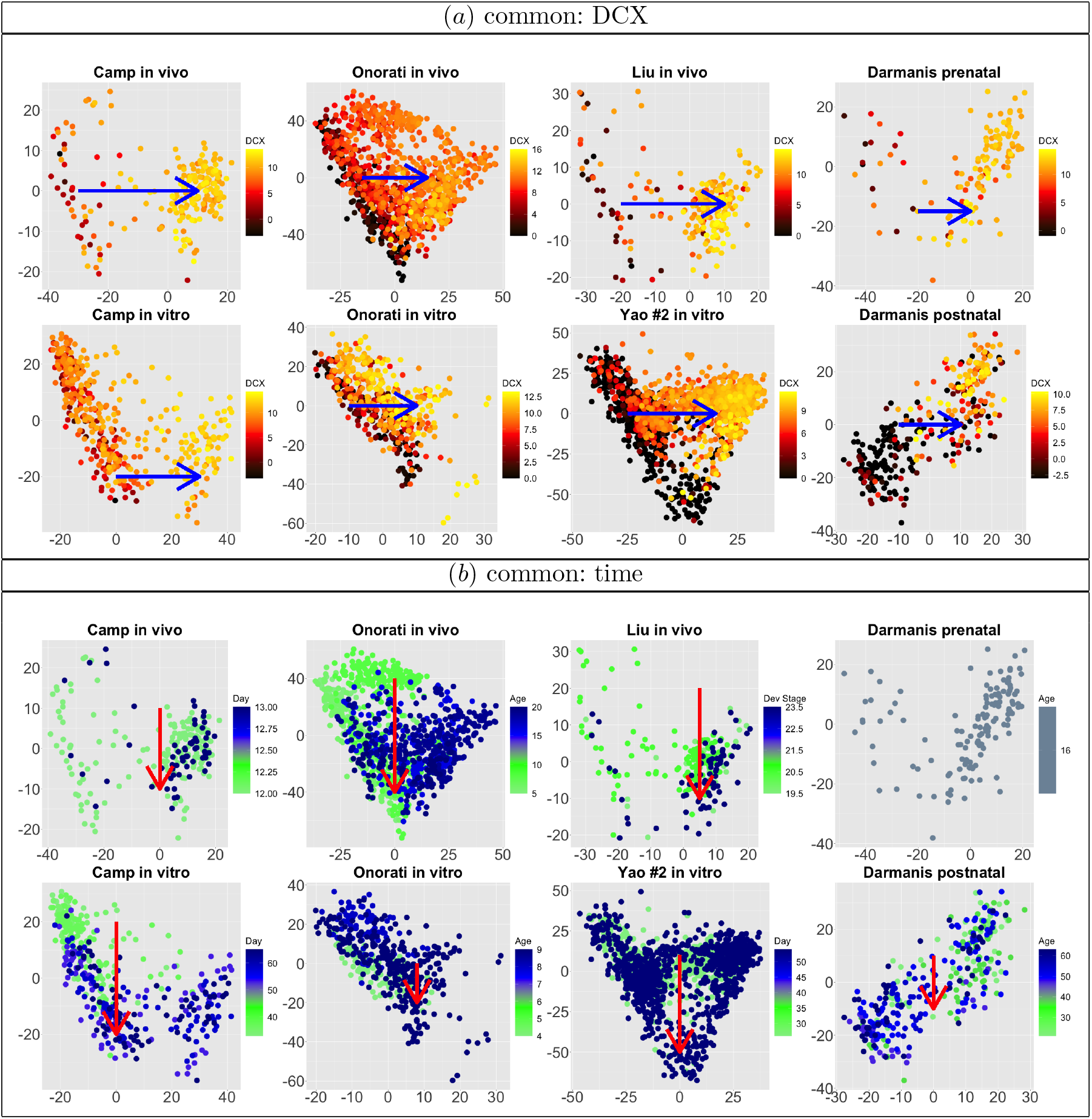
Biological validation by projecting additional data sets onto common components obtained from 2s-LCA. (*a*) Projection of 8 additional scRNA-seq data sets onto the common components with cells colored by the *log*_2_ transformed expression level of the DCX gene. Blue arrows indicate alignment of the first common component with DCX expression (neurogenesis). (*b*) Projection of the same data sets onto the common components with cells colored by time: days of neural differentiation for the *in vitro* data sets and age in years or gestational weeks for the *in vivo* data sets. Red arrows indicate alignment of the second common component with developmental time. The Darmanis *and others* (2015) prenatal study did not specify the exact age of the 4 fetal tissue donors used in their prenatal study, indicating only 16-18 gestational weeks for all samples.

The *in vitro* components generated by 2s-LCA (first 2 panels in the second row of Fig. 3 (*b*) and Fig. 4 (*b*)) were validated by projecting data from another bulk RNA-seq study of neural differentiation onto these components. This replicated the segregation of pluripotent stem cells with high values for the first component, away from other cells in this study that proceeded through neural progenitor and neuronal states, validating the identification of this cell type specific to the *in vitro* studies (see supplementary material, black circles).

## 6. Discussion

In this paper, we proposed two-stage linked component analysis (2s-LCA) for the joint analysis of multiple data sets that are independent but have shared underlying structure resulting from common biological processes and/or shared measurement technologies. The proposed method extracts signal spaces that can be characterized as common, partially shared or individual, which enhances the understanding of the underlying biology between the data sets. Our experimental data results indicate that the 2s-LCA joint decomposition can be a useful tool to define shared molecular dynamics across biologically related data sets, while avoiding unwanted artifacts.

The proposed method remains valid for high dimensional data, as long as the sample covariance matrices are consistent. For the four experimental data sets, we found that the trace and top eigenvalue of the sample covariance matrix for each data set comparable. Hence, it is not unreasonable to assume that the population covariance matrices are of reduced effective ranks, for which the sample covariance matrices are consistent.

The proposed method can be easily extended to high dimensional data sets, where sparsity is necessary or desired. In such cases, the sample covariance matrix estimator used in the proposed method can be replaced by any consistent covariance estimator (e.g., Bickel *and others* (2008); Bien *and others* (2016)).Then the consistency of the proposed 2s-LCA can still be established.

The proposed 2s-LCA depends on a general, but fixed, design. It might also be of interest to extend 2s-LCA to situations without a fixed design or where the design is only partially fixed. There the existence and estimation of subspaces would have to be empirically determined.

## Supporting information

supplemental figures and proofs

## 7. Software

The code to conduct 2s-LCA can be found in the SJD package (https://github.com/CHuanSite/SJD). The four experimental data sets in this paper can be explored at the individual gene level through the NeMO Analytics portal at https://nemoanalytics.org/p?l=ChenEtAl2021&g=DCX.

## 8. Supplementary Material

Supplementary material is available online at https://www.biorxiv.org/

## Acknowledgments

We gratefully acknowledge the comments and suggestions made by the Associate Editor and two referees that led to a much improved paper.

