## supplemental figures and proofs for "Two-stage Linked Component Analysis for Joint Decomposition of Multiple Biologically Related Data Sets"

### S.1 The BEMA Method

**Algorithm 1:** BEMA: standard spiked covariance model space [4]

**Result:** An estimate for the number of spikes, $\hat{K}$

1. $\alpha$ and $\beta$ tuning parameters are set to default value, $\alpha = 0.2, \beta = 0.1$.

2. Write $\tilde{p} = \min(n, p)$. For each $\alpha\tilde{p} \leq k \leq (1-\alpha)\tilde{p}$, obtain $q_k$, the $k/\tilde{p}$-upper-quantile of the MP distribution associated with $\sigma^2 = 1$ and $\gamma_n = p/n$. Compute

$$\hat{\sigma}^2 = \frac{\sum_{\alpha\tilde{p} \leq k \leq (1-\alpha)\tilde{p}} q_k \hat{\lambda}_k}{\sum_{\alpha\tilde{p} \leq k \leq (1-\alpha)\tilde{p}} q_k^2}.$$

3. Obtain $t_{1-\beta}$, the $(1-\beta)$-quantile of Tracy-Widom distribution. Estimate $K$ by

$$\hat{K} = \#\{1 \leq k \leq \tilde{p} : \hat{\lambda}_k > \hat{\sigma}^2[(1+\sqrt{\gamma_n})^2 + t_{1-\beta} \cdot n^{-2/3}\gamma_n^{-1/6}(1+\sqrt{\gamma_n})^{4/3}]\}.$$

Marchenko-Pastur (MP) Distribution: For a given parameter $\gamma > 0$, the Marcenko-Pastur (MP) distribution is defined by the density:

$$f_\gamma(x;\sigma^2) = \frac{1}{2\pi\sigma^2}\frac{1}{x(\gamma \wedge 1)}\sqrt{(x-\sigma^2 h_-)(\sigma^2 h_+ - x)} \cdot 1\{\sigma^2 h- < x < \sigma^2 h_+\},$$

where $h_\pm = (1 \pm \sqrt{\gamma})$. We let $F_\gamma(x;\sigma)$ denote its cumulative distribution function.

Tracy-Widom Distribution: The Tracy-Widom distribution is defined as the limit:

$$F_2(s) = \lim_{n\to\infty} \text{Prob}((\lambda_{\max} - \sqrt{2n})(\sqrt{2n})n^{1/6} \leq s)$$

Where $\lambda_{\max}$ denotes the largest eigenvalue of the random Hermitian matrix.

### S.2 Details on the 2s-LCA Algorithm

**Algorithm 2:** Two-stage linked component analysis for two-way factorial design

**Result:** Estimated signal spaces $\hat{\mathbf{A}}, \hat{\mathbf{B}}_i, \hat{\mathbf{C}}_j, \hat{\mathbf{D}}_{ij}$

1. Extract signal space matrix $\mathbf{U}_{ij}$ for each data set,

$$\hat{\mathbf{U}}_{ij} = \text{top } r_{ij} \text{ eigenvector from } \mathbf{Y}_{ij}\mathbf{Y}_{ij}^T,$$

   determine the $r_{ij}$ by the BEMA method [4].

2. Follow the order

$$\mathbf{A} \rightarrow \mathbf{B}_1 \rightarrow \mathbf{B}_2 \rightarrow \mathbf{C}_1 \rightarrow \mathbf{C}_2 \rightarrow \mathbf{D}_{11} \rightarrow \mathbf{D}_{12} \rightarrow \mathbf{D}_{21} \rightarrow \mathbf{D}_{22}.$$

   to identify the specific signal subspaces as follows.

   (a) Select the number of components (see Algorithm 3 for the general design) and extract principal components from $\hat{\mathbf{U}}_{ij}$ of data sets that share specific signal subspaces;

   (b) Project all $\hat{\mathbf{U}}_{ij}$ onto the space orthogonal to the extracted signal subspaces;

   (c) Repeat the above steps until all signal space has been extracted.

3. Return estimated signal spaces $\hat{\mathbf{A}}, \hat{\mathbf{B}}_i, \hat{\mathbf{C}}_j, \hat{\mathbf{D}}_{ij}$.

**Algorithm 3:** Selection of space dimension $r_k$ for general design

**Result:** An estimator $\hat{r}_k$ for the dimension of the $k$ signal space.

1. Compute $\sum_{i:\Theta_{ik}=1} \hat{\mathbf{U}}_i\hat{\mathbf{U}}_i^T$, and its eigenvalues, $\hat{\sigma}_1, \hat{\sigma}_2, \ldots, \hat{\sigma}_p$.

2. Compute $\eta_1 = \sum_{i:\Theta_{ik}=1} \Theta_{ik}$ and $\eta_2 = \max_{k'} \sum_{i:\Theta_{ik}=1, k' \neq k} \Theta_{ik'}$.

3. Return $\hat{r}_k = \#\{k : \hat{\sigma}_k >= (\eta_1 + \eta_2)/2\}$.

**Algorithm 4:** Two-stage linked component analysis for a general design

**Result:** Estimated signal spaces $\hat{\mathbf{V}}_1, \hat{\mathbf{V}}_2, \ldots, \hat{\mathbf{V}}_K$

1. Extract component space matrix $\hat{\mathbf{U}}_i$ for each data set $i$,

$$\hat{\mathbf{U}}_i = \text{top } \hat{K}_i \text{ eigenvector from } \mathbf{Y}_i \mathbf{Y}_i^T,$$

   where $\hat{K}_i$ is determined by the BEMA method.

2. Follow the order

$$\hat{\mathbf{V}}_1 \to \hat{\mathbf{V}}_2 \cdots \to \hat{\mathbf{V}}_K$$

   to identify the specific components as follows:

   (a) Select the number based on Algorithm 3 and extract principal components from union of data sets that share those components, $\hat{\mathbf{V}}_k$,

$$\text{select } r_k \text{ based on the eigenvalues of } \sum_{i:\Theta_{ik}=1} \hat{\mathbf{U}}_i \hat{\mathbf{U}}_i^T,$$

   and

$$\hat{\mathbf{V}}_k = \text{top } r_k \text{ component from } \sum_{i:\Theta_{ik}=1} \hat{\mathbf{U}}_i \hat{\mathbf{U}}_i^T.$$

   (b) Project the signal space, $\hat{\mathbf{U}}_i$, onto the space orthogonal to the extracted spaces $\hat{\mathbf{V}}_k$,

$$(\mathbf{I} - \hat{\mathbf{V}}_k \hat{\mathbf{V}}_k^T)\hat{\mathbf{U}}_i,$$

   (c) Repeat the above steps until no signal space left.

3. Return estimated signal spaces $\hat{\mathbf{V}}_1, \hat{\mathbf{V}}_2, \ldots, \hat{\mathbf{V}}_K$.

### S.3 Details on Theoretical Results

$$y_{ij} = \Theta_{i1}\mathbf{V}_1 f_{1ij} + \Theta_{i2}\mathbf{V}_2 f_{2ij} + \dots \Theta_{iK}\mathbf{V}_K f_{Kij} + \epsilon_{ij}. \tag{S.1}$$

**Assumption 1.** *$p/n \to \gamma, \gamma > 0$ is a constant.*

**Assumption 2.** *The minimal nonzero eigenvalue of $\mathbf{\Sigma}_{ki}$, denoted by $\sigma_{ki}$, satisfies $\sigma_{ki} \geq \sigma^2(\sqrt{\gamma} + \tau)$, for a positive constant $\tau$ and $\gamma$ in Assumption 1, for all $k$ and $i$.*

**Assumption 3.** *Assume that the random score $f_{kij}$ for each signal subspace $\mathbf{V}_k$ follows a Gaussian distribution, $f_{kij} \sim N(0, \mathbf{\Sigma}_{ki})$, and the noise vector $\epsilon_{ij}$ also follows the Gaussian distribution, $\epsilon_{ij} \sim N(0, \sigma_i^2\mathbf{I})$. as well as the random scores and noises are independent from each other and across subjects.*

**Assumption 4.** *Let $\mathbf{\Sigma}_i = \mathbb{E}[y_{ij}^T y_{ij}]$ for $i = 1, \dots, n$. Assume that these covariance matrices have bounded effective ranks, i.e., $r_e(\mathbf{\Sigma}_i) \leq C$, for all $i$ and some fixed positive constant $C$.*

**Lemma 1** (Davis-Kahan). *[7] Let $\mathbf{\Sigma}, \hat{\mathbf{\Sigma}} \in \mathbb{R}^{p\times p}$ be symmetric, with eigenvalues $\lambda_1 \geq \dots \geq \lambda_p$ and $\hat{\lambda}_1 \geq \dots \geq \hat{\lambda}_p$ respectively. Fix $1 \leq r \leq s \leq p$ and assume that $\min(\lambda_{r-1} - \lambda_r, \lambda_s - \lambda_{s+1}) > 0$, where $\lambda_0 := \infty$ and $\lambda_{p+1} := -\infty$. Let $d := s - r + 1$, and let $\mathbf{V} = (v_r, v_{r+1}, \dots, v_s) \in \mathbb{R}^{p\times d}$ and $\hat{\mathbf{V}} = (\hat{v}_r, \hat{v}_{r+1}, \dots, \hat{v}_s) \in \mathbb{R}^{p\times d}$ have orthonormal columns satisfying $\mathbf{\Sigma} v_j = \lambda_j v_j$ and $\hat{\mathbf{\Sigma}}\hat{v}_j = \hat{\lambda}_j \hat{v}_j$ for $j = r, r+1, \dots, s$. Then*

$$\|\sin\Theta(\hat{\mathbf{V}}, \mathbf{V})\|_F \leq \frac{2\min(d^{1/2}\|\hat{\mathbf{\Sigma}} - \mathbf{\Sigma}\|_{op}, \|\hat{\mathbf{\Sigma}} - \mathbf{\Sigma}\|_F)}{\min(\lambda_{r-1} - \lambda_r, \lambda_s - \lambda_{s+1})}.$$

*Moreover, there exists an orthogonal matrix $\hat{\mathbf{O}} \in \mathbb{R}^{d\times d}$ such that*

$$\|\hat{\mathbf{V}}\hat{\mathbf{O}} - \mathbf{V}\|_F \leq \frac{2^{3/2}\min(d^{1/2}\|\hat{\mathbf{\Sigma}} - \mathbf{\Sigma}\|_{op}, \|\hat{\mathbf{\Sigma}} - \mathbf{\Sigma}\|_F)}{\min(\lambda_{r-1} - \lambda_r, \lambda_s - \lambda_{s+1})}.$$

**Lemma 2.** *Under Model* (S.1)*, Assumptions 1, 2, 4 and $\hat{K}_i$ is from BEMA[**?** ] used in Algorithm 1 with $\alpha \in (0, 1/2)$ a fixed constant and $\beta \to 0$ at a properly rate as $n \to \infty$, then*

$$\hat{K}_i \xrightarrow{P} \sum_{k=1}^{K} \Theta_{ik} r_k.$$

*Proof.* This is a direct application of Lemma 2 in [**?** ]. □

**Lemma 3.** *Let $\hat{\mathbf{\Sigma}}_i$ be the sample covariance matrix for the ith data set, $i = 1, \dots, n$. Under Model* (S.1) *and the assumptions in Lemma 2, we have*

$$\|\hat{\mathbf{\Sigma}}_i - \mathbf{\Sigma}_i\|_F \xrightarrow{P} 0,$$

*and the top $\hat{K}_i$ eigenvectors $\hat{\mathbf{U}}_i$ of $\hat{\mathbf{\Sigma}}_i$ converge in probability to the top $\sum_{k=1}^{K}\Theta_{ik}r_k$ eigenvectors $\mathbf{U}_i$ of population covariance matrix $\mathbf{\Sigma}_i$, i.e.,*

$$\hat{\mathbf{U}}_i \xrightarrow[\text{in space}]{P} \mathbf{U}_i.$$

*Proof.* By Theorem 2.1 in [1], there exists a constant $c_1$ such that

$$\|\hat{\mathbf{\Sigma}}_i - \mathbf{\Sigma}_i\|_F \leq 2c_1 \cdot \|\mathbf{\Sigma}_i\|_2 \cdot r_e(\mathbf{\Sigma}_i) \cdot \sqrt{\frac{\ln n}{n}},$$

with probability $1 - 5n^{-1}$. Thus, when $r_e(\mathbf{\Sigma}_i) \leq C$, $\|\hat{\mathbf{\Sigma}}_i - \mathbf{\Sigma}_i\|_F \xrightarrow{P} 0$. Then, by Davis-Kahan theorem,

$$\hat{\mathbf{U}}_i \xrightarrow[\text{in space}]{P} \mathbf{U}_i,$$

which completes the proof. □

**Lemma 4.** *Under Model S.1, Assumption 1, 2, 4, and based on Lemma 2, 3, we have for any k,*

$$\sum_{i:\Theta_{ik}=1} \hat{\mathbf{U}}_i \hat{\mathbf{U}}_i^T \xrightarrow{P} \sum_{i:\Theta_{ik}=1} \mathbf{U}_i \mathbf{U}_i^T.$$

*Proof.* To prove it, based on the Davis-Kahan Theorem 1, there exists one rotation matrix $O$, such that

$$\begin{aligned}
\|\mathbf{U}_i\mathbf{U}_i^T - \hat{\mathbf{U}}_\mathbf{i}\hat{\mathbf{U}}_{\mathbf{i}}{}_i^T\|_F &= \|\mathbf{U}_i\mathbf{U_i}^T - \hat{\mathbf{U}}_\mathbf{i} OO^T \hat{\mathbf{U}}_i^T\|_F \\
&= \|\mathbf{U}_i\mathbf{U}_i^T - \hat{\mathbf{U}}_i OO^T \hat{\mathbf{U}}_i^T\|_F \\
&= \|\mathbf{U}_i\mathbf{U}_i^T - \mathbf{U}_i O\hat{\mathbf{U}}_i^T + \mathbf{U}_i O\hat{\mathbf{U}}_i^T + \hat{\mathbf{U}}_i OO^T \hat{\mathbf{U}}_i^T\|_F \\
&\leq \|\mathbf{U}_i\|_F \|\mathbf{U_i}^T - O\hat{\mathbf{U}}_i^T\|_F + \|\mathbf{U}_i O - \hat{\mathbf{U}}_\mathbf{i}\|_F \|O^T \hat{\mathbf{U}}_i\|_F.
\end{aligned}$$

According to [7],

$$\|\mathbf{U}_i\mathbf{U}_i^T - \hat{\mathbf{U}}_i\hat{\mathbf{U}}_i^T\|_F \xrightarrow{P} 0.$$

when $\hat{\Sigma}_i \xrightarrow{P} \Sigma_i$ in Frobenius Norm. This also indicates that

$$\sum_{i:\Theta_{ik}=1} \hat{\mathbf{U}}_i \hat{\mathbf{U}}_i^T \xrightarrow{P} \sum_{i:\Theta_{ik}=1} \mathbf{U}_i \mathbf{U}_i^T.$$

in the Frobenius norm. □

**Lemma 5.** *Under Model S.1, Assumption 1, 2, 4 and based on Lemma 2, 3, 4, by choosing the eigenvectors* $\hat{\mathbf{V}}_k$ *of* $\sum_{i,\Theta_{ik}=1} \hat{\mathbf{U}}_i \hat{\mathbf{U}}_i^T$ *with eigenvalues larger than* $(\eta_1 + \eta_2)/2$*, where* $\eta_1 = \sum_{i,\Theta_{ik}=1} \Theta_{ik}$ *and* $\eta_2 = \max_{k'} \sum_{i:\Theta_{ik}=1, k \neq k} \Theta_{ik'}$*, then*

$$\hat{\mathbf{V}}_k \xrightarrow[\textit{in space}]{P} \mathbf{V}_k$$

*Proof.* Note that

$$\sum_{i:\Theta_{ik}=1} \hat{\mathbf{U}}_i \hat{\mathbf{U}}_i^T \xrightarrow{P} \sum_{i:\Theta_{ik}=1} \mathbf{U}_i \mathbf{U}_i^T.$$

Then based on the continuity of the eigenvalue operator, we have that

$$\hat{r}_k = \text{Number of eigenvalues of} \sum_{i:\Theta_{ik}=1} \hat{\mathbf{U}}_i \hat{\mathbf{U}}_i^T \text{ larger than } (\eta_1 + \eta_2)/2$$

$\hat{r}_k$ is consistent for $r_k$, so that

$$\hat{\mathbf{V}}_k \xrightarrow[\text{in space}]{P} \mathbf{V}_k.$$

□

**Lemma 6.** *Under Model S.1, the Lemma 1, 2, 4, and based on Lemma 2, 3, 4, 5, we have that*

$$(\mathbf{I} - \hat{\mathbf{V}}_k \hat{\mathbf{V}}_k^t)\hat{\mathbf{U}}_i \xrightarrow[\textit{in space}]{P} (\mathbf{I} - \mathbf{V}_k \mathbf{V}_k^t)\mathbf{U}_i.$$

*Proof.* Note that in Lemma 5, we have that

$$\hat{\mathbf{V}}_k \xrightarrow[\text{in space}]{P} \mathbf{V}_k,$$

and in Lemma 4, we have that

$$\hat{\mathbf{U}}_k \xrightarrow[\text{in space}]{P} \mathbf{U}_k,$$

Then, according to the Continuous Mapping Theorem [3],

$$(\mathbf{I} - \hat{\mathbf{V}}_k \hat{\mathbf{V}}_k^t)\hat{\mathbf{U}}_i \xrightarrow[\text{in space}]{P} (\mathbf{I} - \mathbf{V}_k \mathbf{V}_k^t)\mathbf{U}_i.$$

holds. □

**Theorem 1.** *Under Model* (S.1)*, if Assumptions 1 - 4 hold, then*

$$\hat{\mathbf{V}}_k \xrightarrow[\textit{in space}]{P} \mathbf{V}_k, \textit{ for all } k.$$

*Proof.* By recursively applying Lemmas 2, 3, and 4, 5, 6 in the appendix, consistency is proved. □

### S.4 Additional Simulation on Existing Method Comparisons

#### S.4.1 Fix rank

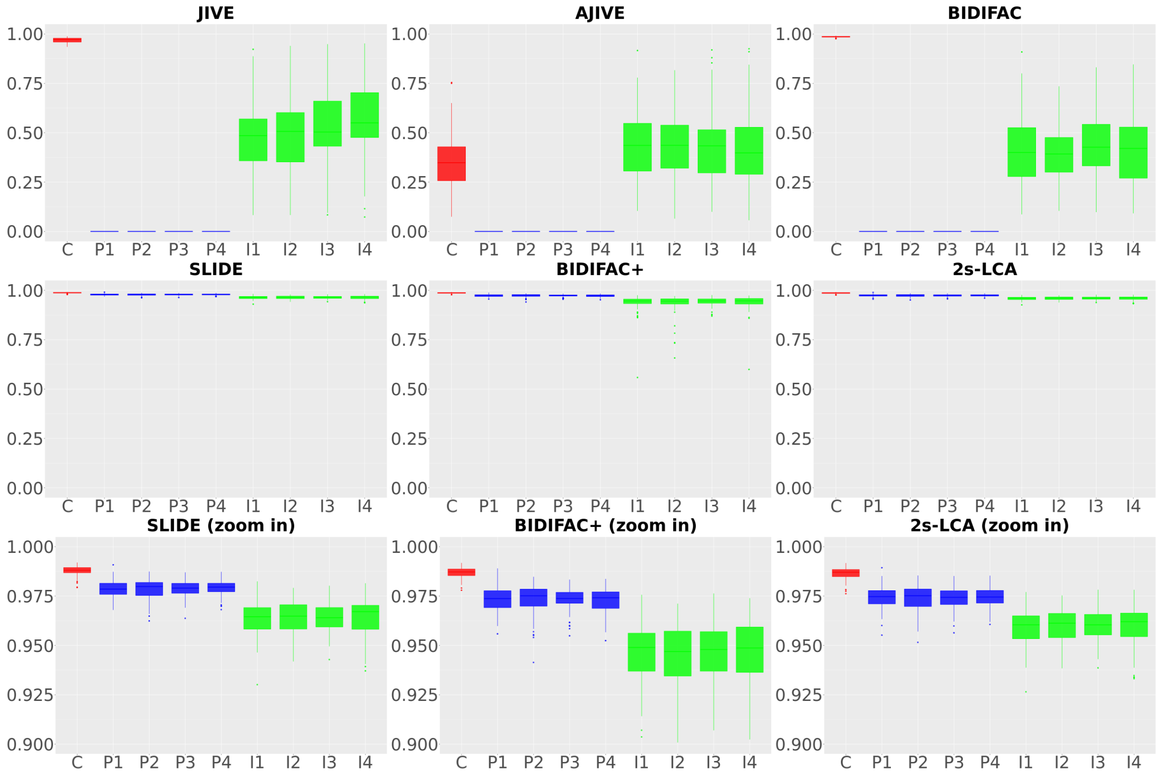


Figure S.1: Comparisons between JIVE, AJIVE, SLIDE, BIDIFAC, BIDIFAC+ and 2s-LCA. A total number of 1000 simulations are run for each method under the low dimensional setting, where $p = 50$, $n = 500$, SNR $= 0.2$.

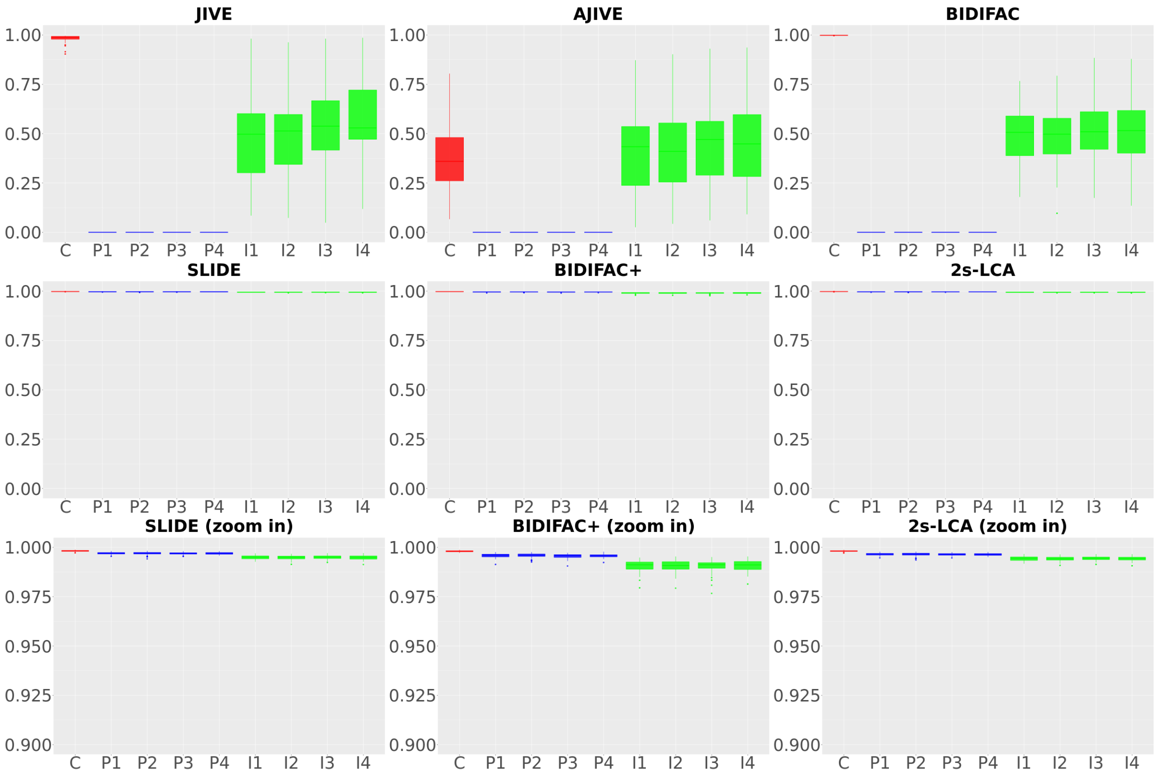


Figure S.2: Comparisons between JIVE, AJIVE, SLIDE, BIDIFAC, BIDIFAC+ and 2s-LCA. A total number of 1000 simulations are run for each method under the low dimensional setting, where $p = 50$, $n = 500$, SNR $= 1$.

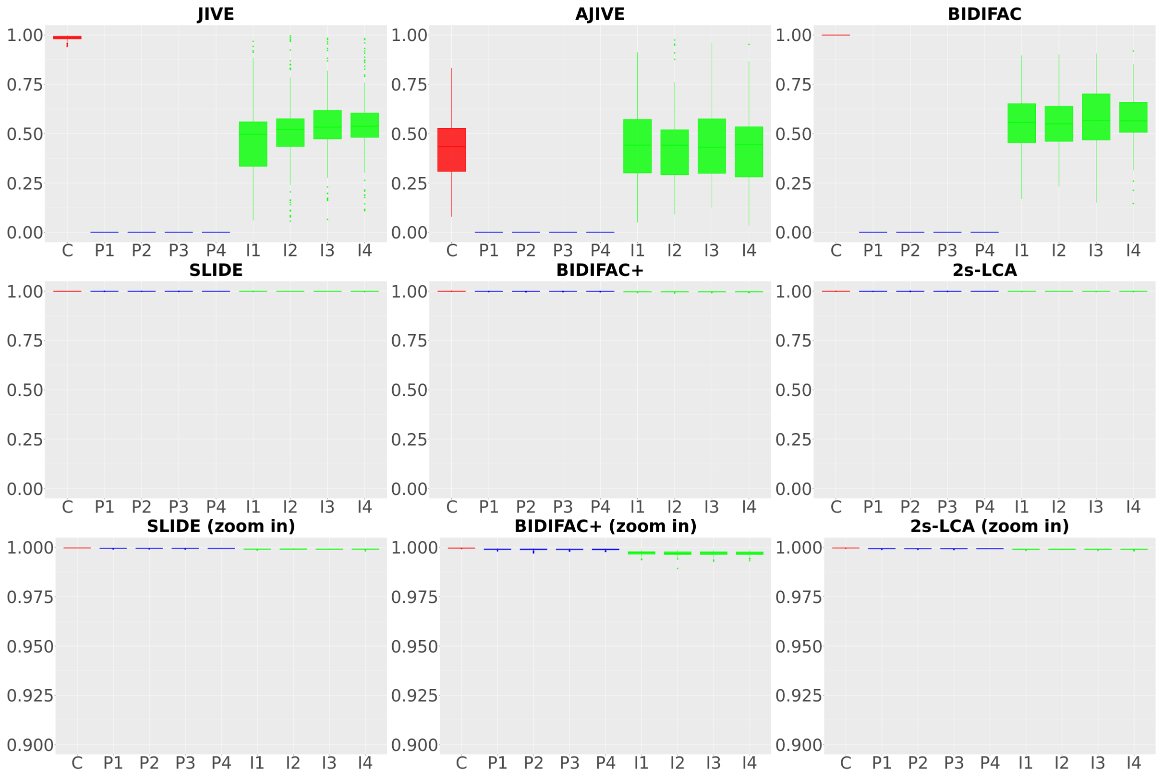


Figure S.3: Comparisons between JIVE, AJIVE, SLIDE, BIDIFAC, BIDIFAC+ and 2s-LCA. A total number of 1000 simulations are run for each method under the low dimensional setting, where $p = 50$, $n = 500$, SNR $= 5$.

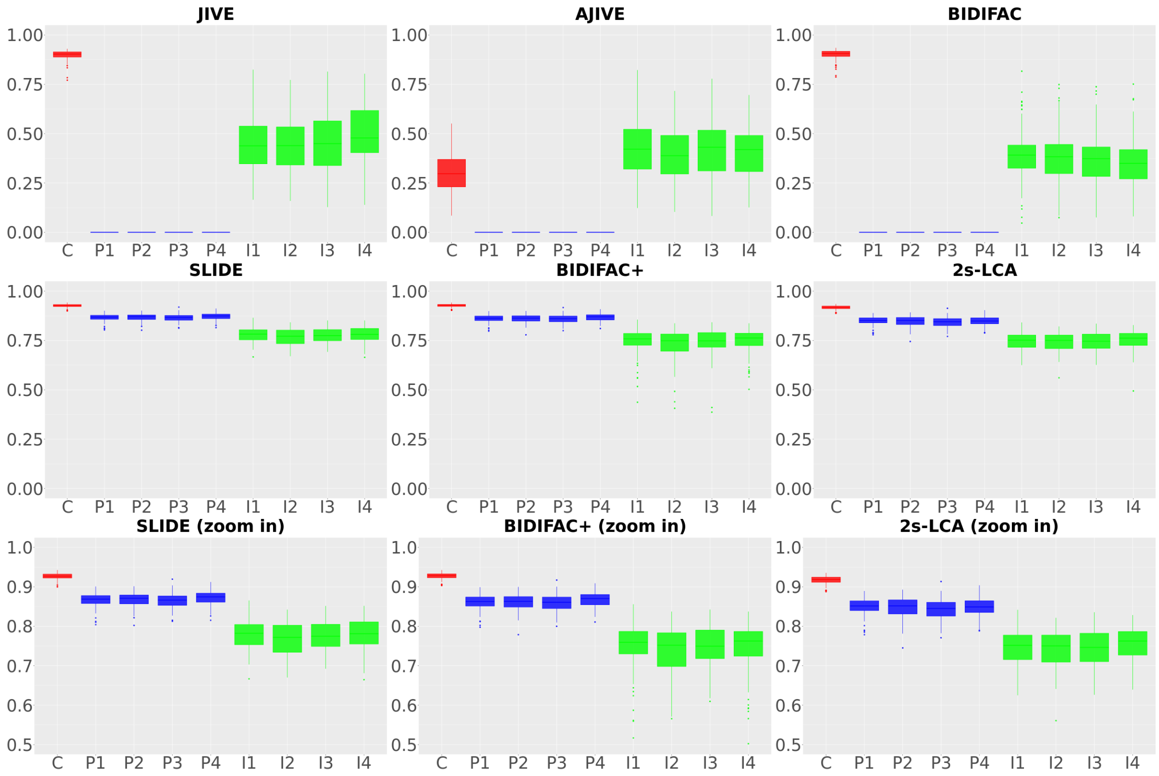


Figure S.4: Comparisons between JIVE, AJIVE, SLIDE, BIDIFAC, BIDIFAC+ and 2s-LCA. A total number of $1000$ simulations are run for each method under the low dimensional setting, where $p = 500$, $n = 50$, SNR $= 0.2$.

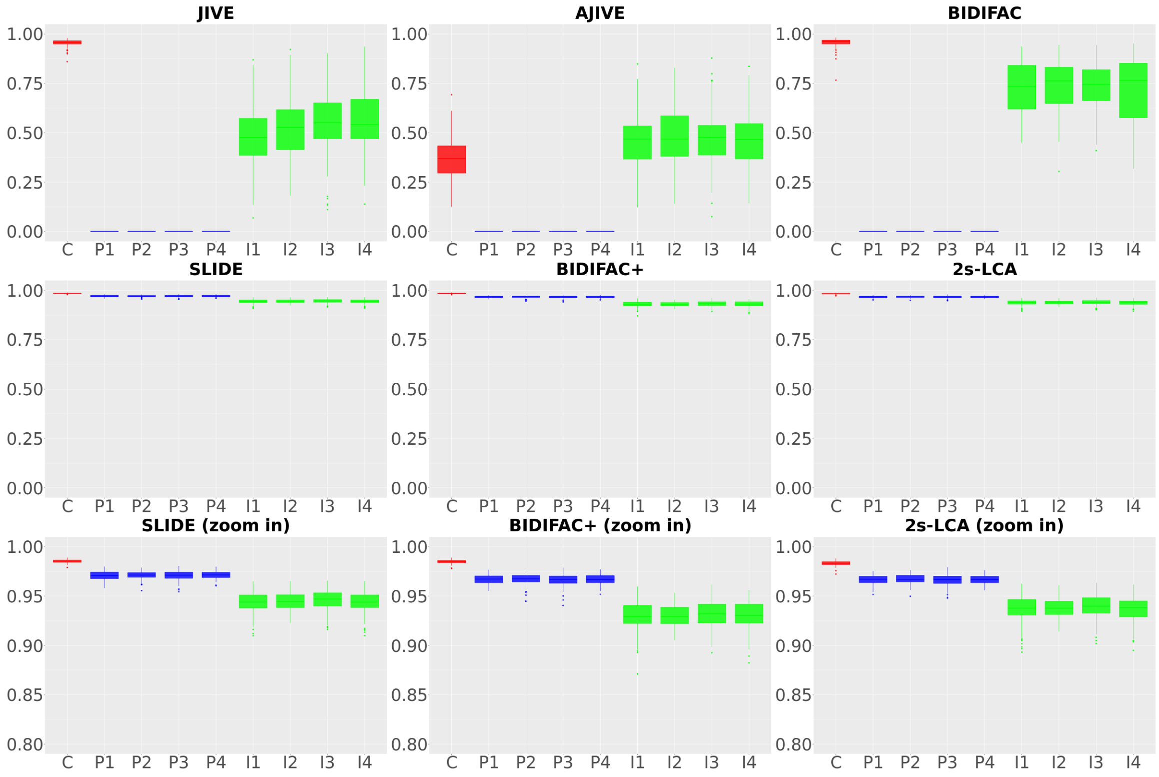


Figure S.5: Comparisons between JIVE, AJIVE, SLIDE, BIDIFAC, BIDIFAC+ and 2s-LCA. A total number of 1000 simulations are run for each method under the low dimensional setting, where $p = 500$, $n = 50$, SNR $= 1$.

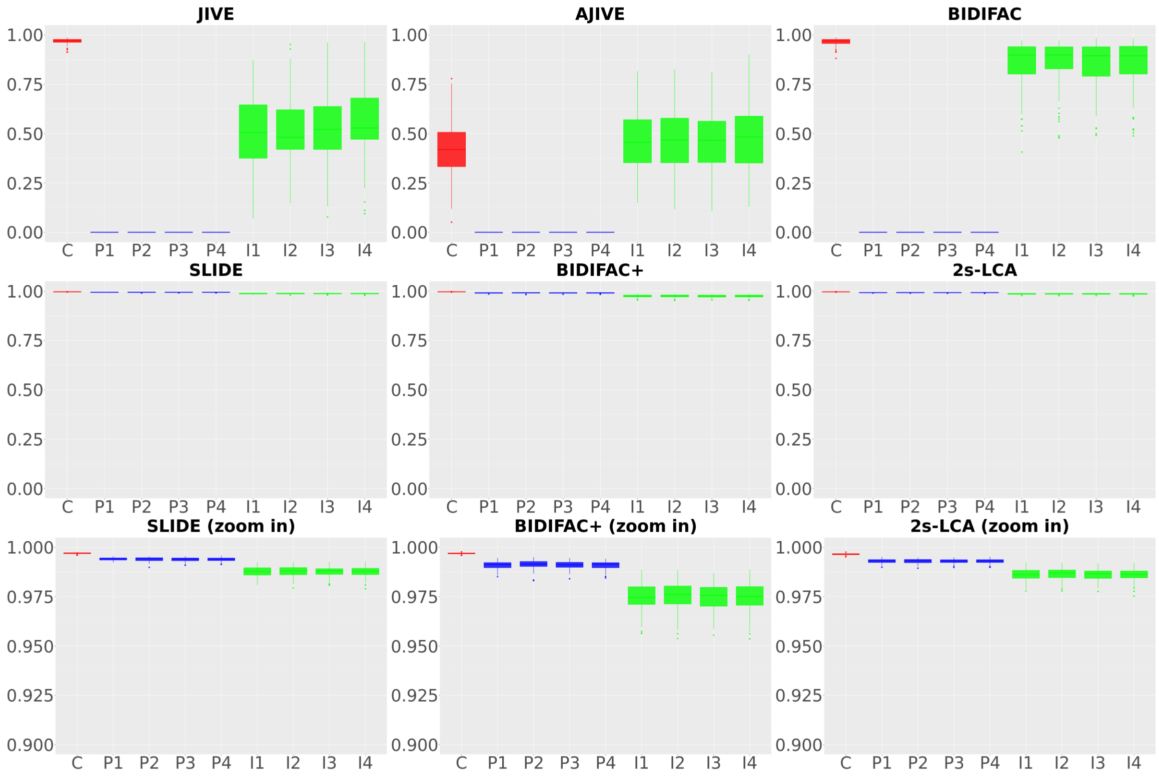


Figure S.6: Comparisons between JIVE, AJIVE, SLIDE, BIDIFAC, BIDIFAC+ and 2s-LCA. A total number of 1000 simulations are run for each method under the low dimensional setting, where $p = 500$, $n = 50$, SNR $= 5$.

#### S.4.2 Different Individual Component Variation

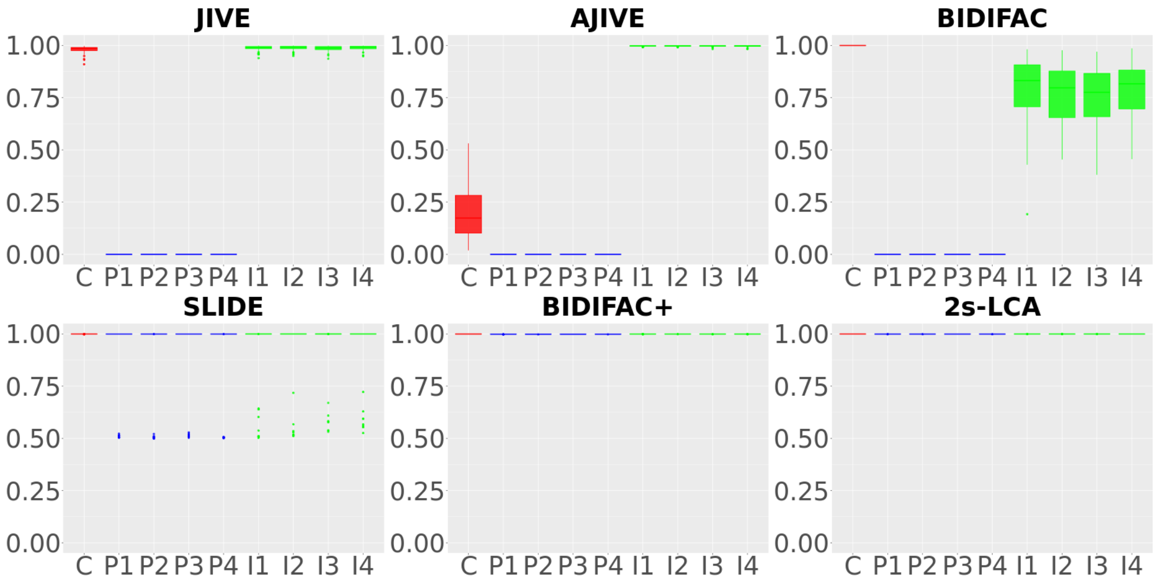


Figure S.7: Comparisons between JIVE, AJIVE, SLIDE, BIDIFAC, BIDIFAC+ and 2s-LCA. A total number of 1000 simulations are run for each method under the low dimensional setting, where $p = 50$, $n = 500$, ivsf $= 10$, here ivsf means "individual variance scaling factor", which means that we multiply the individual component's variance by this factor in the simulated data.

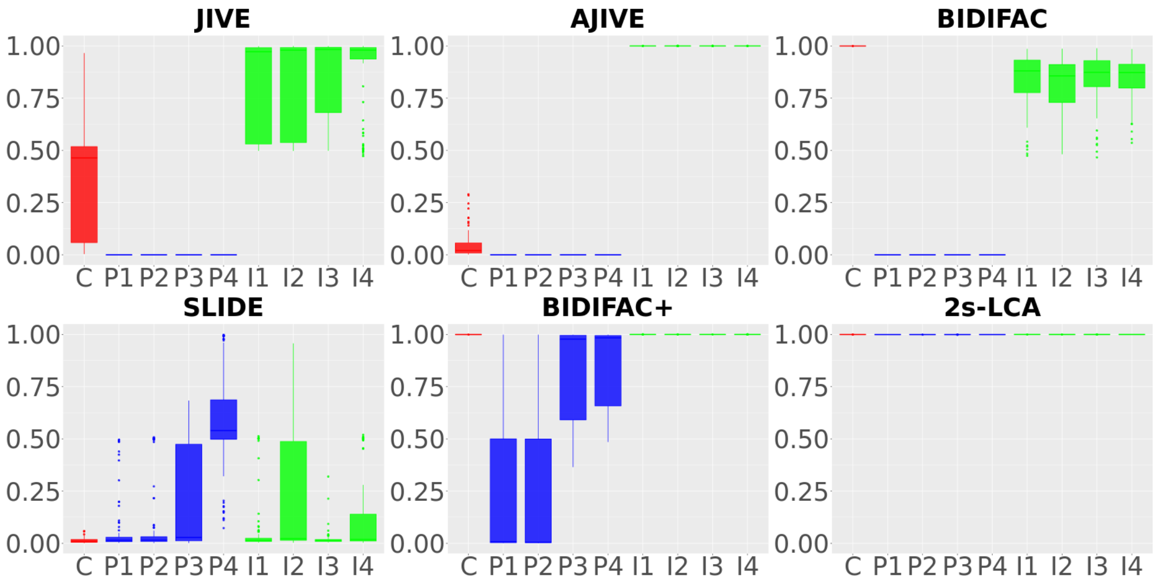


Figure S.8: Comparisons between JIVE, AJIVE, SLIDE, BIDIFAC, BIDIFAC+ and 2s-LCA. A total number of 1000 simulations are run for each method under the low dimensional setting, where $p = 50$, $n = 500$, ivsf $= 1000$, here ivsf means "individual variance scaling factor", which means that we multiply the individual component's variance by this factor in the simulated data.

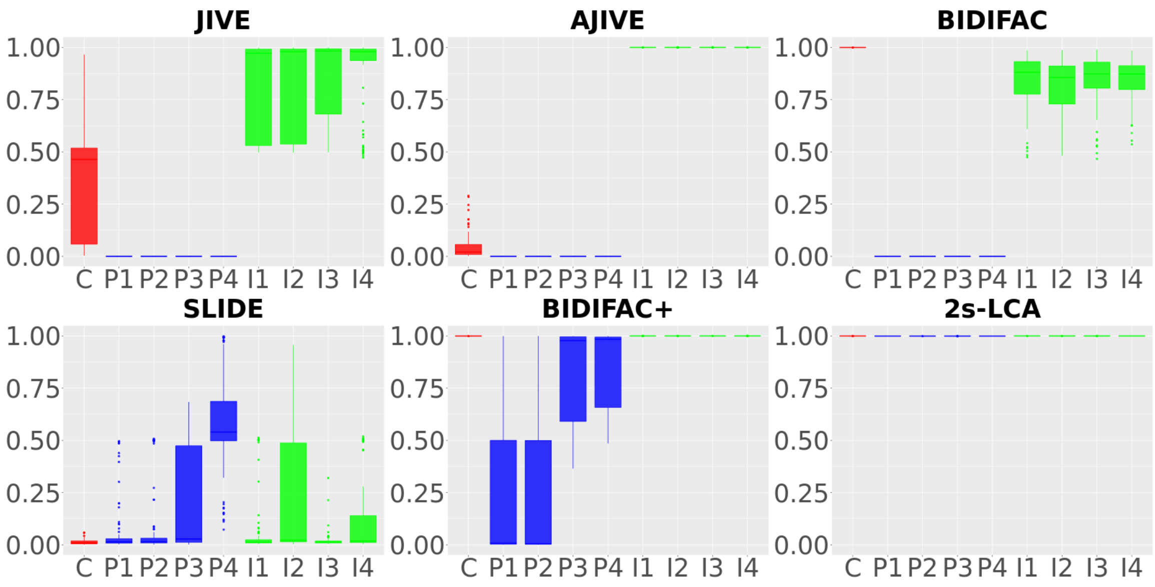


Figure S.9: Comparisons between JIVE, AJIVE, SLIDE, BIDIFAC, BIDIFAC+ and 2s-LCA. A total number of 1000 simulations are run for each method under the low dimensional setting, where $p = 50$, $n = 500$, ivsf $= 100000$, here ivsf means "individual variance scaling factor", which means that we multiply the individual component's variance by this factor in the simulated data.

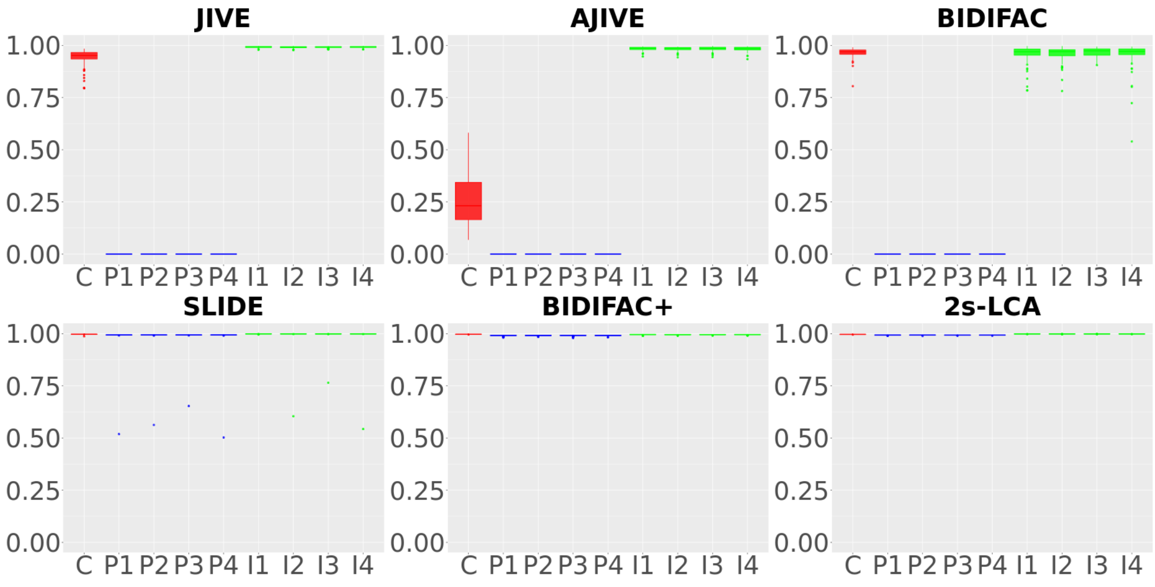


Figure S.10: Comparisons between JIVE, AJIVE, SLIDE, BIDIFAC, BIDIFAC+ and 2s-LCA. A total number of 1000 simulations are run for each method under the low dimensional setting, where $p = 500$, $n = 50$, ivsf $= 10$, here ivsf means "individual variance scaling factor", which means that we multiply the individual component's variance by this factor in the simulated data.

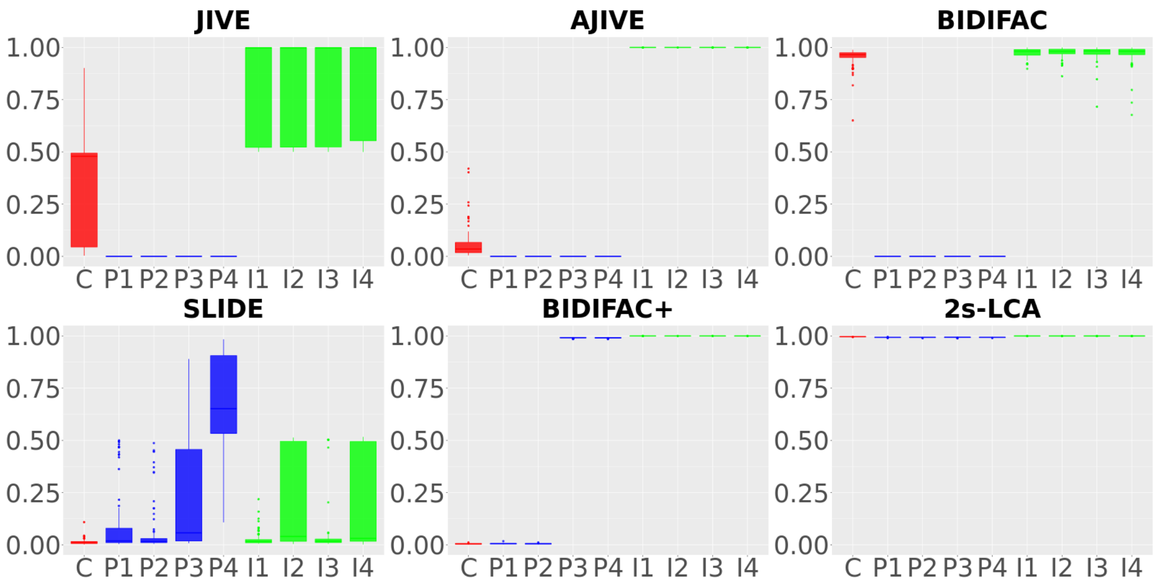


Figure S.11: Comparisons between JIVE, AJIVE, SLIDE, BIDIFAC, BIDIFAC+ and 2s-LCA. A total number of 1000 simulations are run for each method under the low dimensional setting, where $p = 500$, $n = 50$, ivsf $= 1000$, here ivsf means ”individual variance scaling factor”, which means that we multiply the individual component’s variance by this factor in the simulated data.

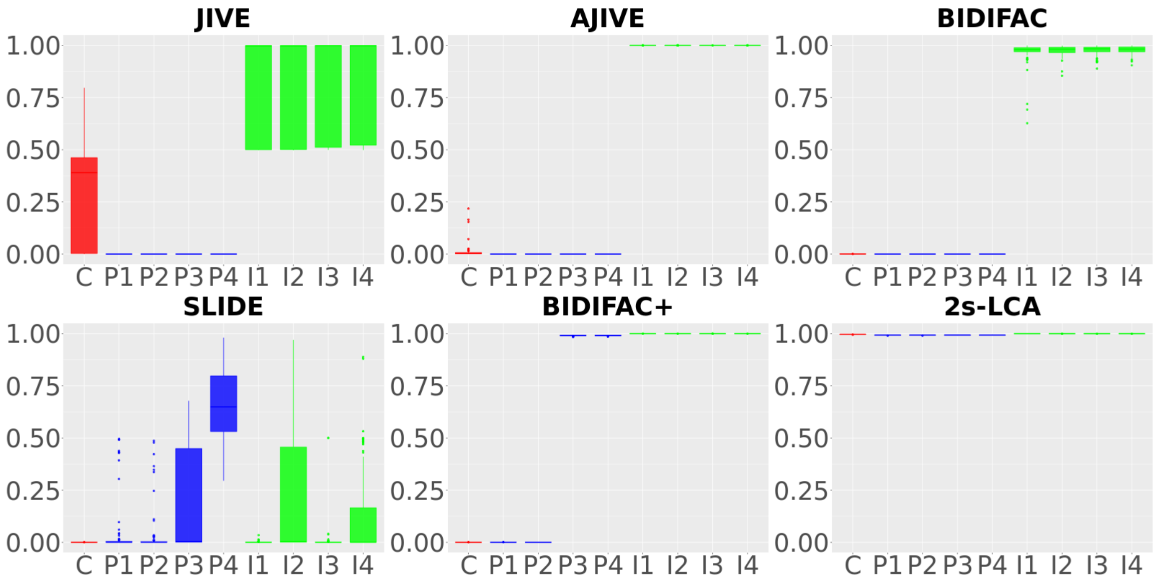


Figure S.12: Comparisons between JIVE, AJIVE, SLIDE, BIDIFAC, BIDIFAC+ and 2s-LCA. A total number of 1000 simulations are run for each method under the low dimensional setting, where $p = 500$, $n = 50$, ivsf $= 100000$, here ivsf means ”individual variance scaling factor”, which means that we multiply the individual component’s variance by this factor in the simulated data.

### S.4.3 Rank Selection

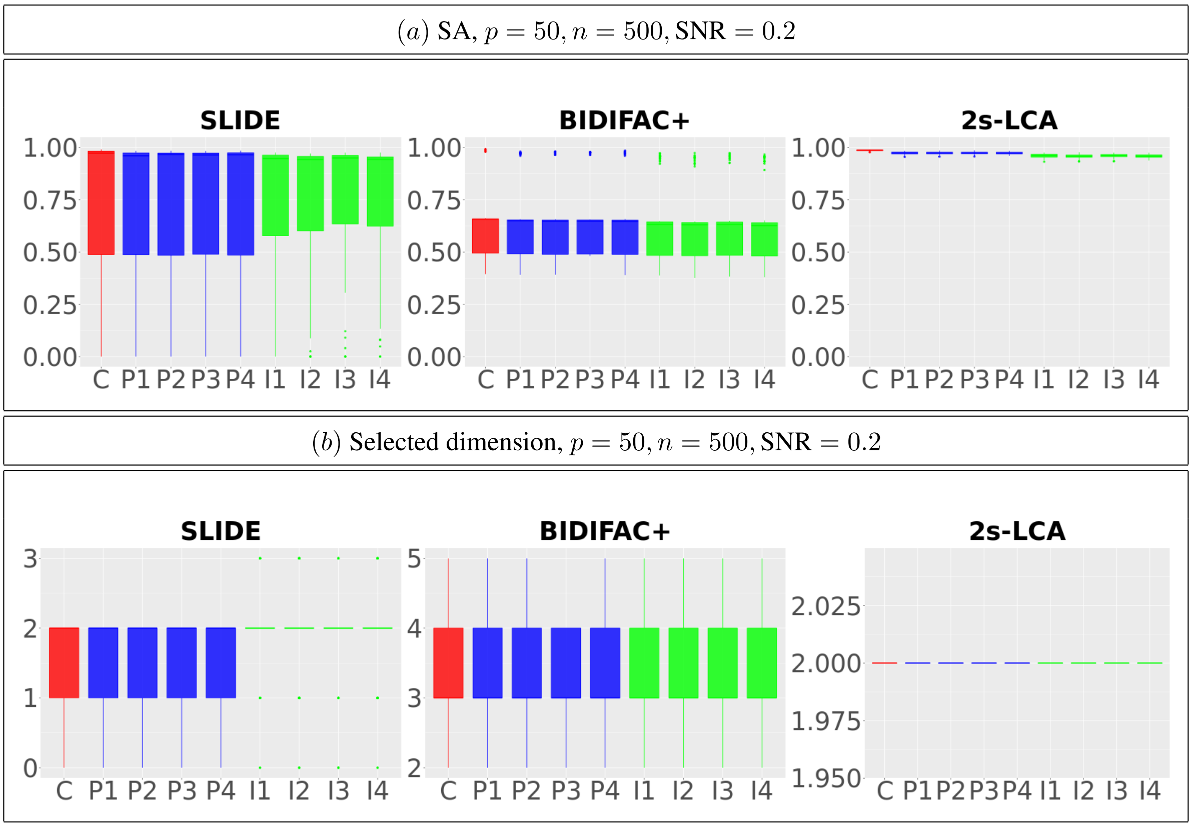


Figure S.13: Comparisons between SLIDE, BIDIFAC+ and 2s-LCA. $(a)$ Boxplot for PVE; $(b)$ Boxplot for rank selection. A total number of 1000 simulations are run for each method under the low dimensional setting, where $p = 50$ and $n = 500$ and SNR $= 0.2$.

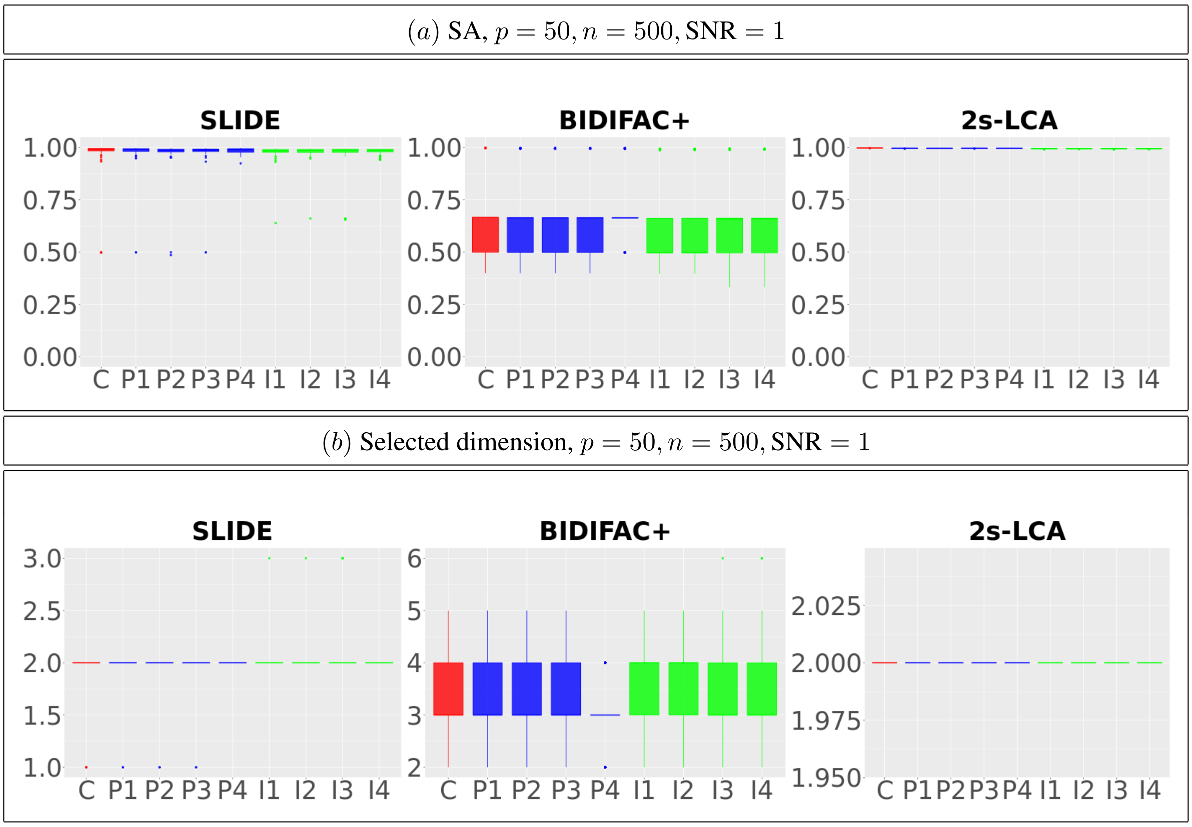


Figure S.14: Comparisons between SLIDE, BIDIFAC+ and 2s-LCA. $(a)$ Boxplot for SA; $(b)$ Boxplot for rank selection. A total number of $1000$ simulations are run for each method under the low dimensional setting, where $p = 50$ and $n = 500$ and $\text{SNR} = 1$.

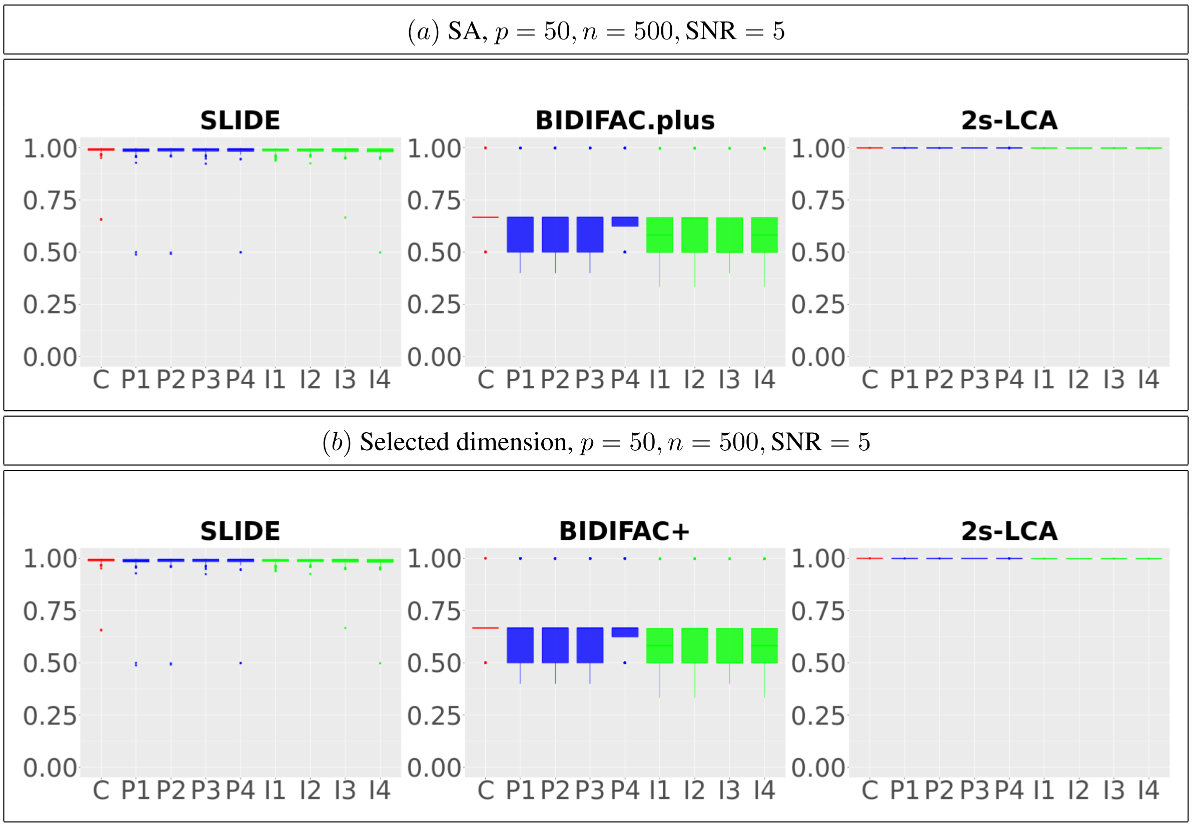


Figure S.15: Comparisons between SLIDE, BIDIFAC+ and 2s-LCA. $(a)$ Boxplot for SA; $(b)$ Boxplot for rank selection. A total number of $1000$ simulations are run for each method under the low dimensional setting, where $p = 50$ and $n = 500$ and $\mathrm{SNR} = 5$.

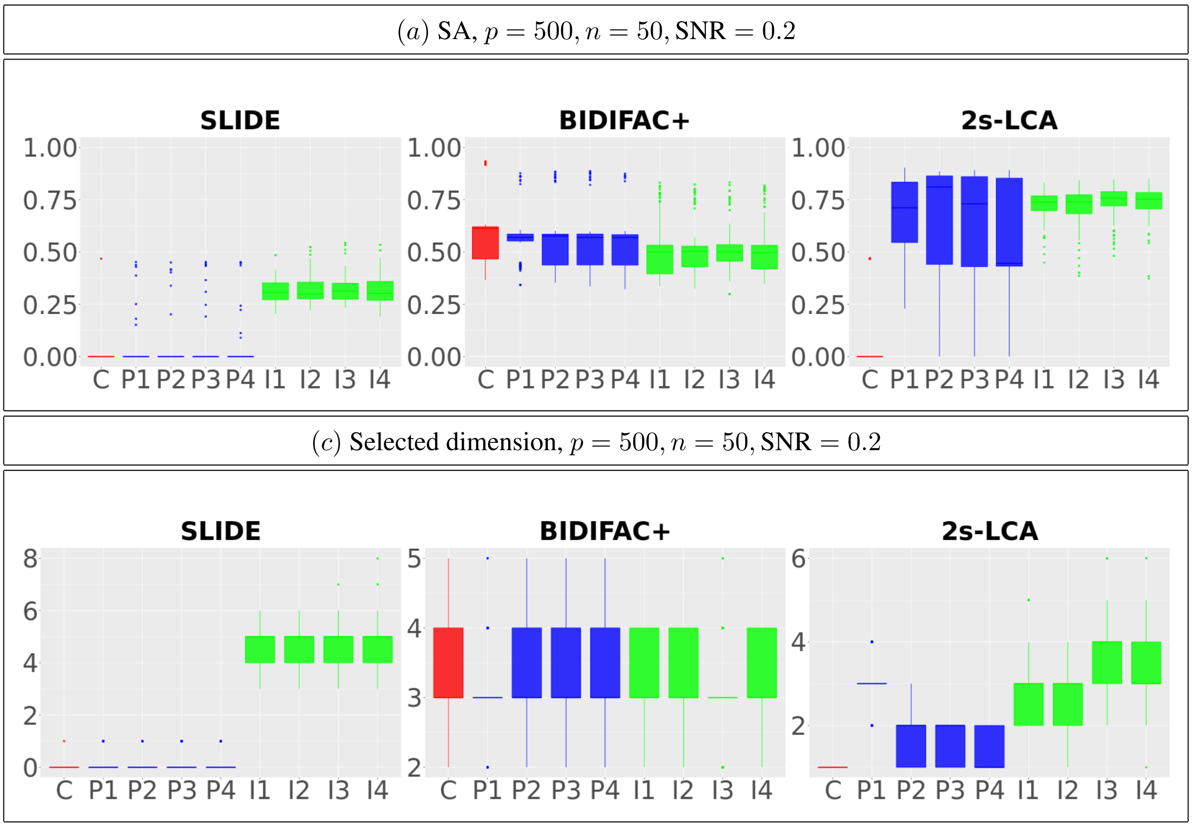


Figure S.16: Comparisons between SLIDE, BIDIFAC+ and 2s-LCA. $(a)$ Boxplot for SA; $(b)$ Boxplot for rank selection. A total number of 1000 simulations are run for each method under the high dimensional setting, where $p = 500$ and $n = 50$ and $\text{SNR} = 0.2$.

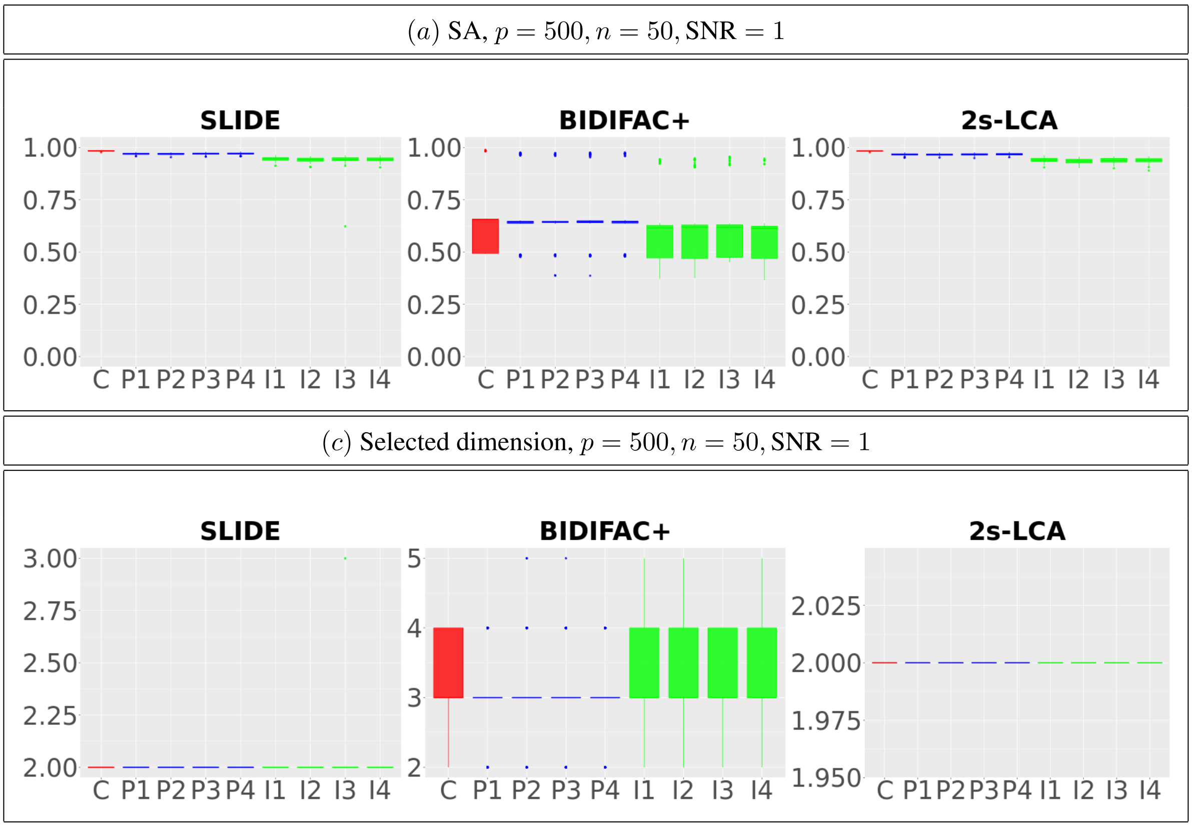


Figure S.17: Comparisons between SLIDE, BIDIFAC+ and 2s-LCA. ($a$) Boxplot for SA; ($b$) Boxplot for rank selection. A total number of 1000 simulations are run for each method under the high dimensional setting, where $p = 500$ and $n = 50$ and SNR $= 1$.

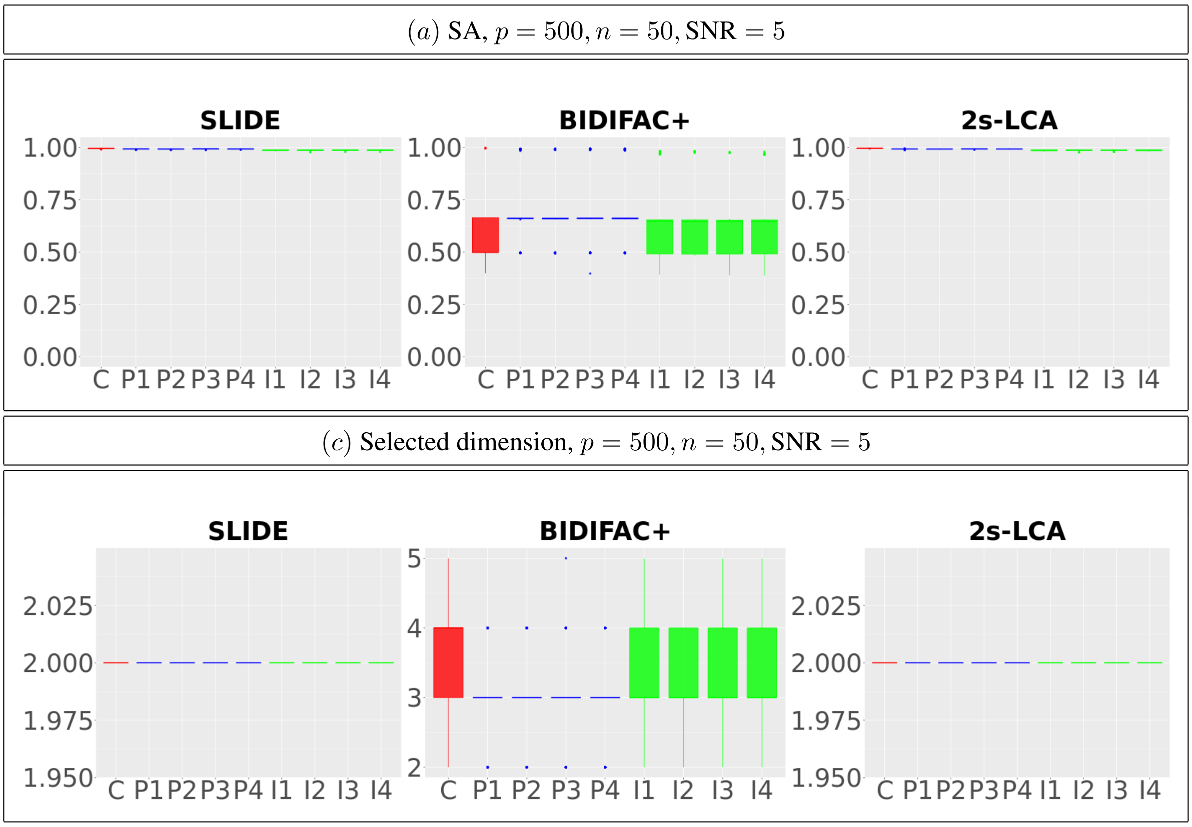


Figure S.18: Comparisons between SLIDE, BIDIFAC+ and 2s-LCA. $(a)$ Boxplot for SA; $(b)$ Boxplot for rank selection. A total number of 1000 simulations are run for each method under the high dimensional setting, where $p = 500$ and $n = 50$ and SNR $= 5$.

### S.5 Additional Analysis of Gene Loadings

#### S.5.1 Loadings for genes in 1st common component

Table S.1: Top genes with its dcx value and time

Table S.2: Top 1 - 25 genes

| Gene | 1-dcx | 2-time |
|---|---|---|
| STMN2 | 0.047804535 | 0.091587513 |
| RTN1 | 0.040272709 | 0.056599444 |
| CHL1 | 0.034247689 | 0.031053639 |
| MYT1L | 0.031277677 | 0.051534255 |
| INA | 0.028930952 | 0.055559835 |
| GRIA2 | 0.028137561 | 0.03263489 |
| NRXN1 | 0.027829555 | 0.030200783 |
| SCN3B | 0.027813977 | 0.031207797 |
| DCX | 0.027322386 | 0.061143006 |
| MEF2C | 0.026683239 | 0.025640036 |
| CXCL14 | 0.026510667 | -0.023417776 |
| SCN3A | 0.026425857 | 0.026417441 |
| GAP43 | 0.025848973 | 0.046918435 |
| NTRK2 | 0.025764666 | -0.0366064 |
| STMN4 | 0.024646616 | 0.016268088 |
| SCN2A | 0.023821944 | 0.022723925 |
| KIF5A | 0.023435139 | 0.031042151 |
| PCSK1N | 0.023219871 | 0.002804714 |
| CAMK2B | 0.02315452 | 0.006517724 |
| DLG2 | 0.023026609 | 0.025997539 |
| MIAT | 0.022916118 | 0.02491282 |
| ATP1B2 | 0.022904298 | -0.033853999 |
| CELF4 | 0.022529811 | 0.027077858 |
| GPM6A | 0.022040541 | 0.027845968 |
| ERBB4 | 0.02193002 | 0.005768301 |

Table S.3: Top 26 - 50 genes

| Gene | 1-dcx | 2-time |
|---|---|---|
| ANK2 | 0.021885749 | 0.026593107 |
| RUNX1T1 | 0.021669083 | 0.047308243 |
| MALAT1 | 0.021651183 | -0.016602158 |
| NCAM1 | 0.021639913 | 0.037832003 |
| ARPP21 | 0.021524596 | 0.022826531 |
| SOBP | 0.021488185 | 0.025568619 |
| KIDINS220 | 0.021462819 | 0.036983722 |
| TTC3 | 0.021348765 | 0.018562876 |
| ANK3 | 0.021190775 | 0.036192019 |
| NDRG4 | 0.021045609 | 0.005234878 |
| GNAO1 | 0.020803216 | 0.015582532 |
| APC2 | 0.020554546 | 0.016615135 |
| NTM | 0.02054747 | 0.005600546 |
| SLIT1 | 0.020471567 | 0.024940109 |
| SYT1 | 0.020471075 | 0.048580028 |
| SPARCL1 | 0.020458447 | -0.037396392 |
| CADPS | 0.020373134 | 0.023417602 |
| PSD2 | 0.020117152 | 0.005673743 |
| MAP1B | 0.020023364 | 0.030963851 |
| TSPAN7 | 0.019930451 | 0.000522618 |
| PCDH9 | 0.019657476 | -0.002685756 |
| SCD5 | 0.019454492 | 0.014558608 |
| AKAP9 | 0.019253839 | 0.027805073 |
| SNAP25 | 0.019249166 | 0.032721988 |
| SST | 0.019242449 | 0.000129336 |

#### S.5.2 Loadings for genes in *in vitro* components

Table S.4: Top genes expression values

Table S.5: Top 1 - 25 genes

| Gene | 1-pluripotency | 2 |
|---|---|---|
| TDGF1 | 0.046391026 | -0.025086856 |
| POU5F1 | 0.045500092 | -0.015831052 |
| GAL | 0.038854842 | 0.002245821 |
| CER1 | 0.033345693 | -0.000196294 |
| PODXL | 0.03266612 | 0.003164171 |
| HHLA1 | 0.031125068 | -0.016128503 |
| PRDM14 | 0.0306378 | -0.002842124 |
| CDH1 | 0.030024799 | 0.010020738 |
| GABRB3 | 0.029934717 | -0.005183345 |
| SFPQ | 0.029498573 | 0.000507492 |
| NOP56 | 0.029300006 | -0.009504269 |
| IDO1 | 0.028771741 | -0.011122691 |
| S100A10 | 0.028760278 | 0.010018084 |
| FLT1 | 0.028470098 | 0.008184392 |
| TNFRSF8 | 0.028318038 | -0.011297958 |
| PDPN | 0.028074427 | -0.005719546 |
| TUBA4A | 0.027950545 | -0.006822115 |
| LCK | 0.027929881 | -0.010282376 |
| LEFTY1 | 0.027927035 | -0.015717727 |
| SCG3 | 0.027893703 | 0.00121996 |
| HTR7 | 0.027383897 | -0.000305728 |
| SLC22A3 | 0.027284478 | 0.000584988 |
| CLDN7 | 0.027004383 | -0.000106976 |
| POLR3G | 0.026804085 | -0.004860002 |
| GJA4 | 0.026751055 | 0.006876659 |

Table S.6: Top 26 - 50 genes

| Gene | 1-pluripotency | 2 |
|---|---|---|
| CAPN13 | 0.026704465 | 0.005198376 |
| SLC7A5 | 0.026660591 | -0.001884795 |
| NETO1 | 0.026626432 | 0.015131885 |
| VRTN | 0.026540478 | -0.01015922 |
| PYCARD | 0.026537153 | -0.008628315 |
| TMSB4X | 0.026466363 | 0.006571814 |
| SOX14 | 0.026381279 | 0.006548762 |
| FAM83F | 0.026346208 | -0.00496873 |
| TUBB2A | 0.026220354 | -0.008125069 |
| SYK | 0.026123832 | 0.014988279 |
| GABRP | 0.026067299 | 0.004245249 |
| ACTA1 | 0.025867972 | 0.048043765 |
| ZFP42 | 0.025733962 | 0.013568464 |
| GRID2 | 0.025724491 | 0.010489039 |
| SCGB3A2 | 0.025722441 | 0.001123915 |
| INA | 0.025612181 | 0.0209179 |
| LGALS12 | 0.025366754 | 0.001180065 |
| ZDHHC22 | 0.025279054 | -0.00308465 |
| TUBA1C | 0.025197237 | -0.02842435 |
| RAB17 | 0.025156635 | -0.00536432 |
| NANOG | 0.02490072 | -0.013019514 |
| MGST1 | 0.02476074 | 8.03E-05 |
| HSPA8 | 0.024753183 | -0.010628604 |
| EIF5A | 0.024737705 | 0.001174462 |
| CHGA | 0.024708481 | 0.005462141 |

### S.6 Additional Results on Change Order of 2s-LCA

#### S.6.1 bulk-sc-invitro-invivo

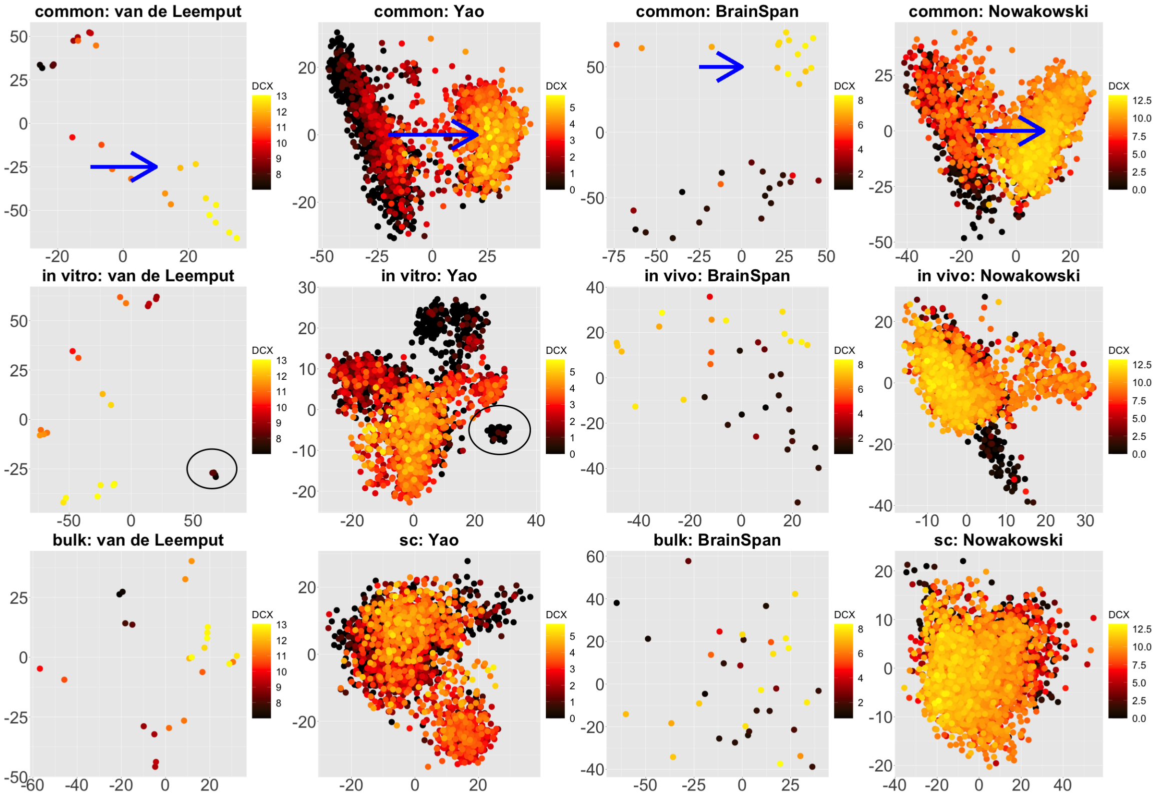


Figure S.19: Scatterplots of scores of each data set corresponding to common components (top panel), partially shared components associated with environments (middle panel: *in vitro* on the left two plots and *in vivo* on the right two plots), and partially shared components associated with technologies (bottom panel: bulk on the left two plots and single cell on the right two plots) by 2s-LCA.

#### S.6.2 bulk-sc-invivo-invitro

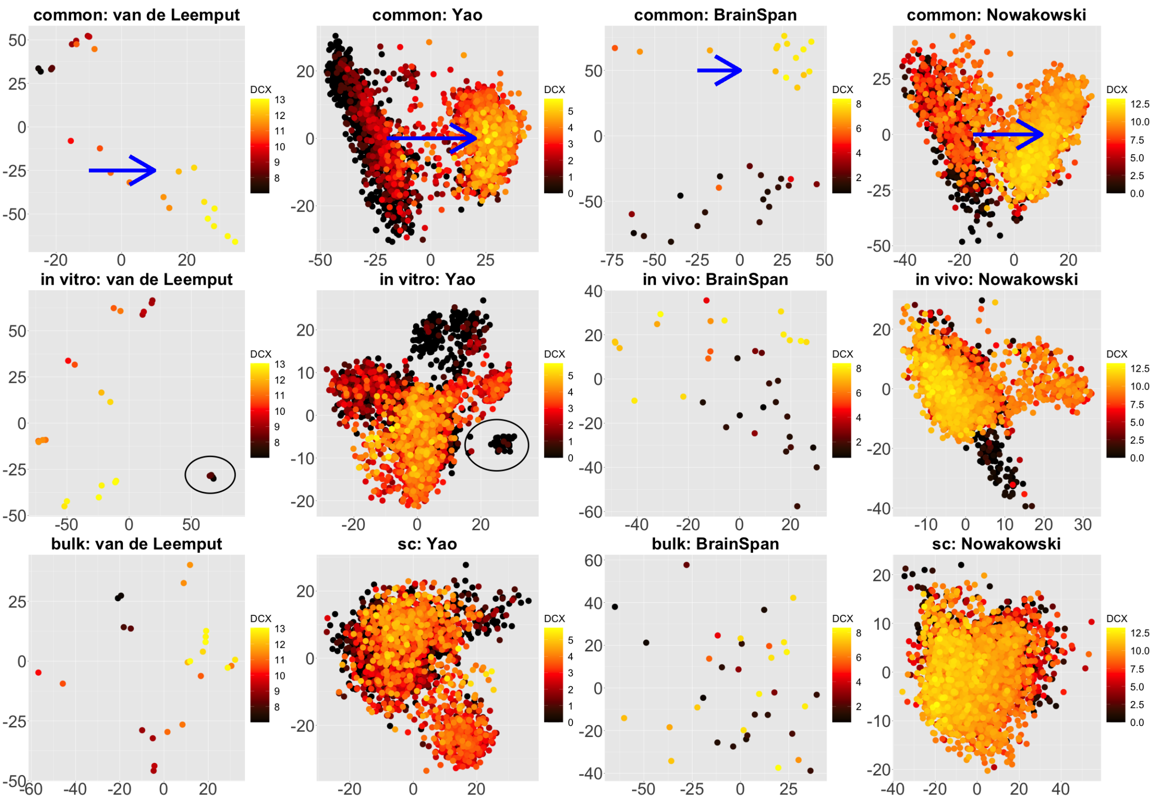


Figure S.20: Scatterplots of scores of each data set corresponding to common components (top panel), partially shared components associated with environments (middle panel: *in vitro* on the left two plots and *in vivo* on the right two plots), and partially shared components associated with technologies (bottom panel: bulk on the left two plots and single cell on the right two plots) by 2s-LCA.

### S.6.3 invitro-invivo-bulk-sc

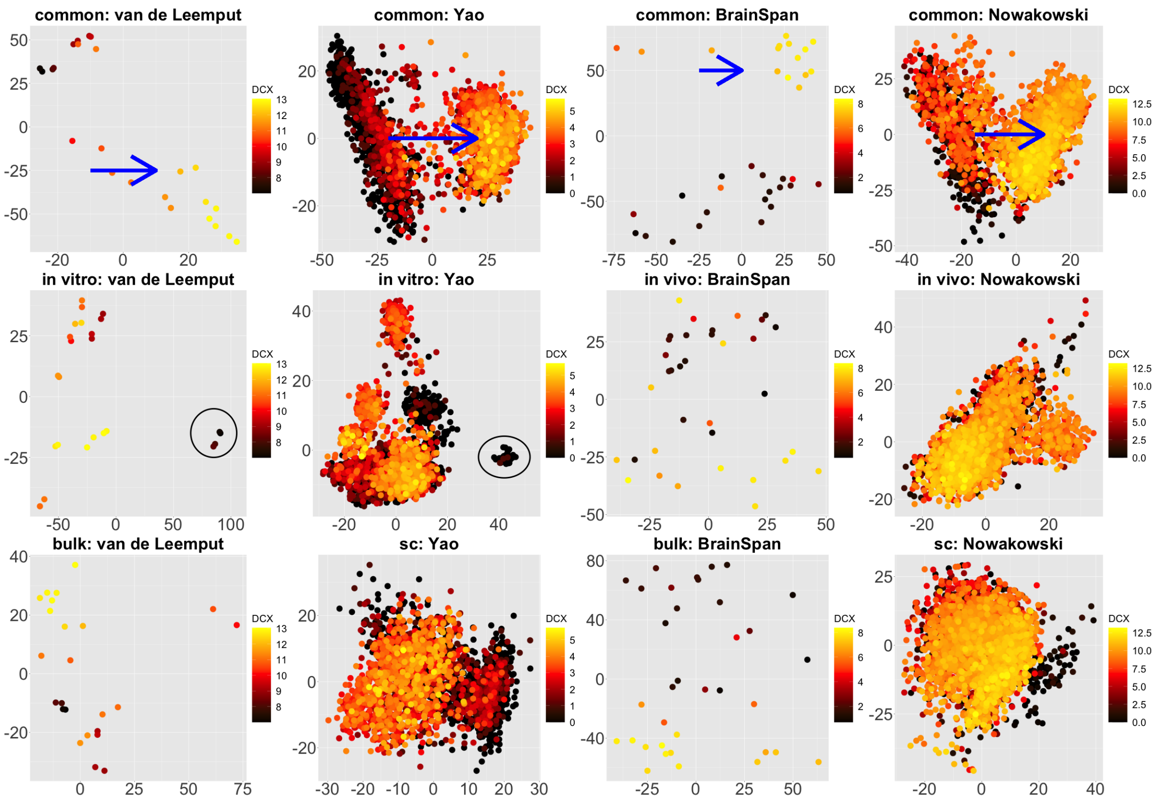


Figure S.21: Scatterplots of scores of each data set corresponding to common components (top panel), partially shared components associated with environments (middle panel: *in vitro* on the left two plots and *in vivo* on the right two plots), and partially shared components associated with technologies (bottom panel: bulk on the left two plots and single cell on the right two plots) by 2s-LCA.

#### S.6.4 invitro-invivo-sc-bulk

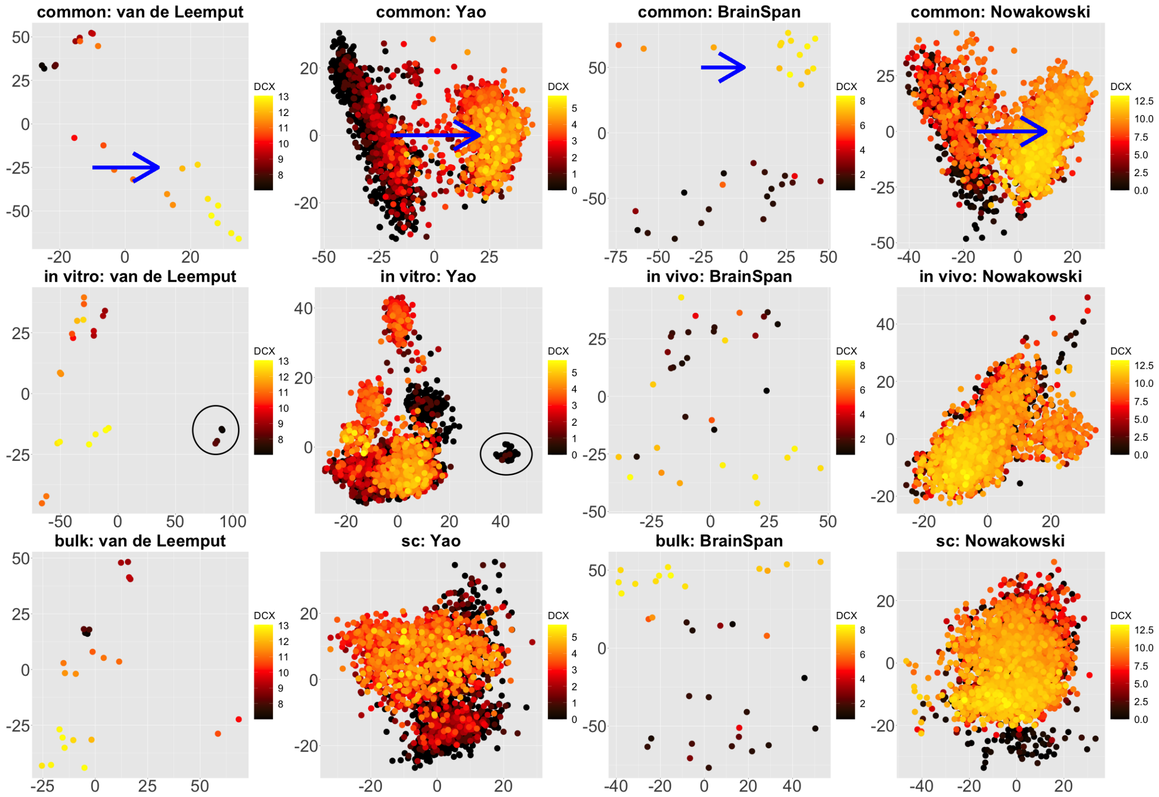


Figure S.22: Scatterplots of scores of each data set corresponding to common components (top panel), partially shared components associated with environments (middle panel: *in vitro* on the left two plots and *in vivo* on the right two plots), and partially shared components associated with technologies (bottom panel: bulk on the left two plots and single cell on the right two plots) by 2s-LCA.

#### S.6.5 invivo-invitro-bulk-sc

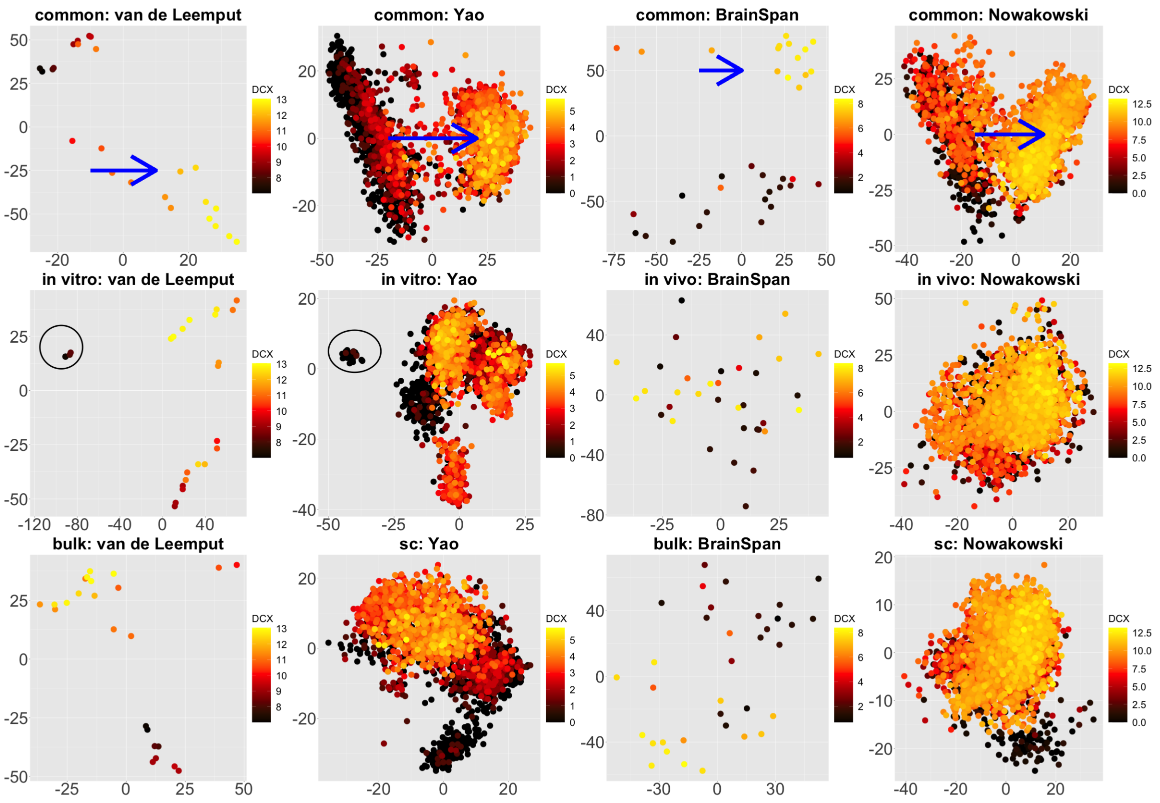


Figure S.23: Scatterplots of scores of each data set corresponding to common components (top panel), partially shared components associated with environments (middle panel: *in vitro* on the left two plots and *in vivo* on the right two plots), and partially shared components associated with technologies (bottom panel: bulk on the left two plots and single cell on the right two plots) by 2s-LCA.

#### S.6.6 invivo-invitro-sc-bulk

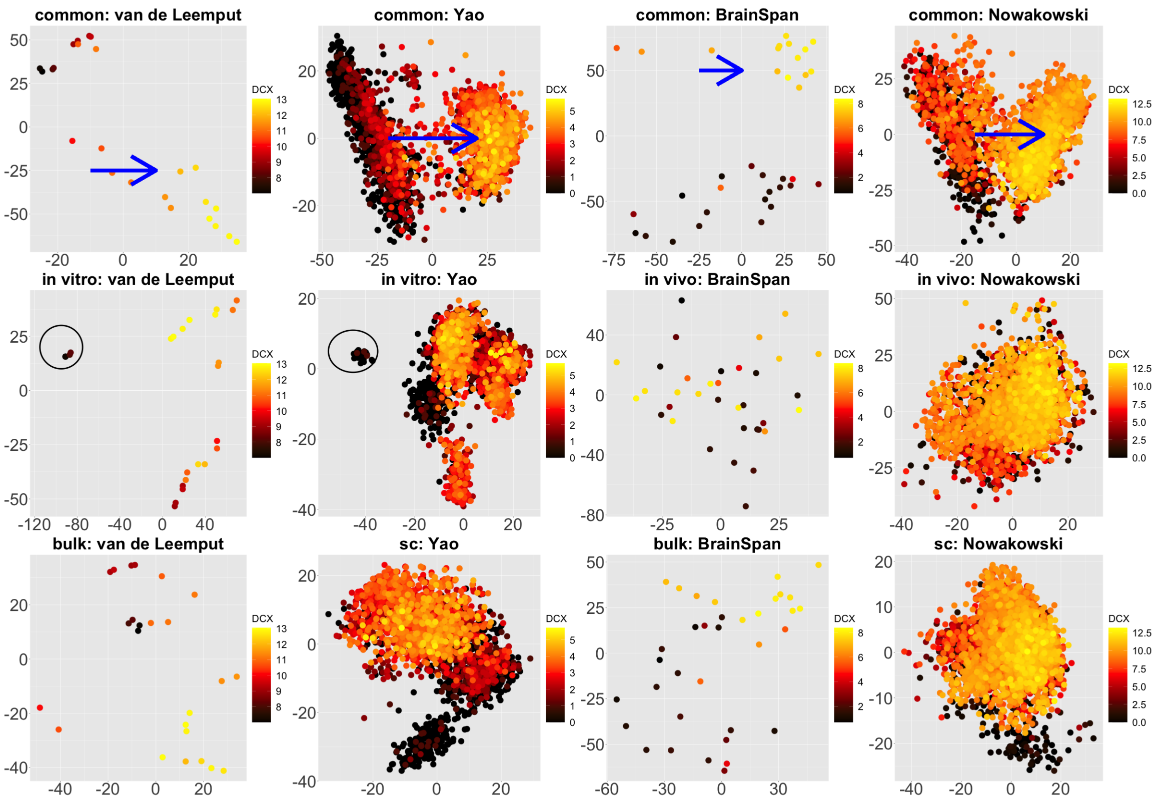


Figure S.24: Scatterplots of scores of each data set corresponding to common components (top panel), partially shared components associated with environments (middle panel: *in vitro* on the left two plots and *in vivo* on the right two plots), and partially shared components associated with technologies (bottom panel: bulk on the left two plots and single cell on the right two plots) by 2s-LCA.

#### S.6.7 sc-bulk-invitro-invivo

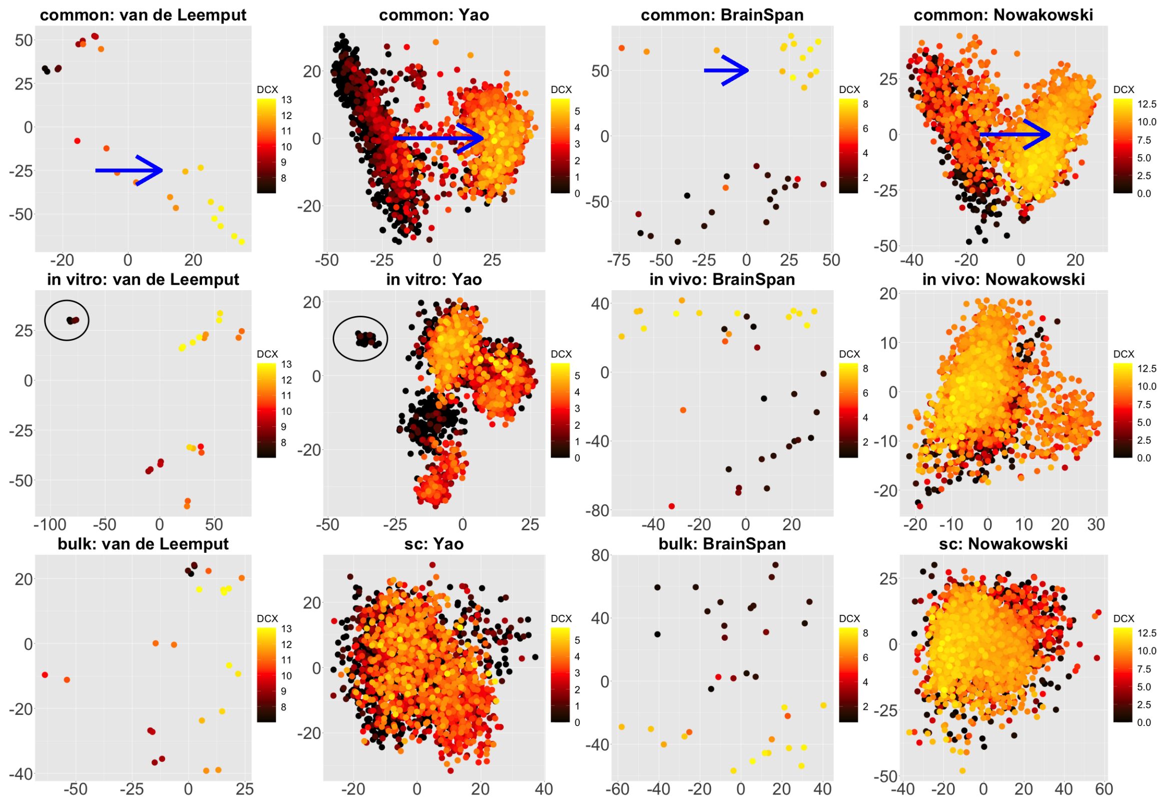


Figure S.25: Scatterplots of scores of each data set corresponding to common components (top panel), partially shared components associated with environments (middle panel: *in vitro* on the left two plots and *in vivo* on the right two plots), and partially shared components associated with technologies (bottom panel: bulk on the left two plots and single cell on the right two plots) by 2s-LCA.

### S.6.8 sc-bulk-invivo-invitro

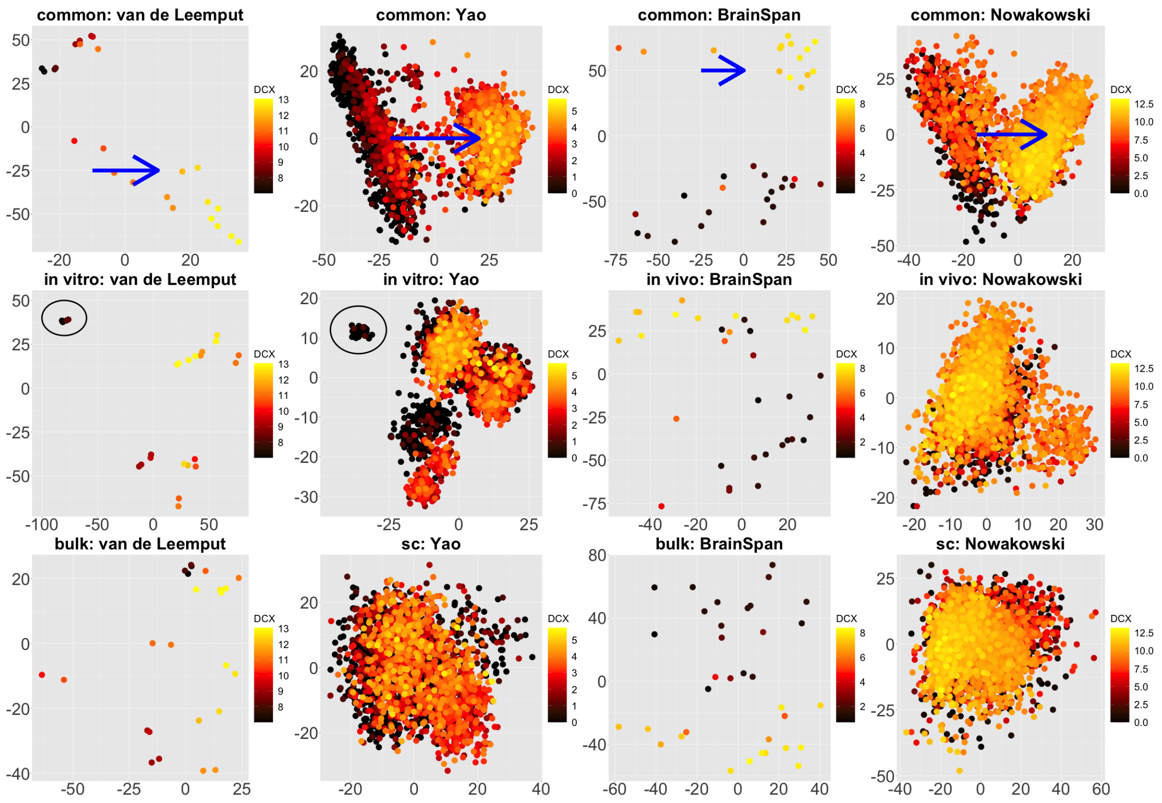


Figure S.26: Scatterplots of scores of each data set corresponding to common components (top panel), partially shared components associated with environments (middle panel: *in vitro* on the left two plots and *in vivo* on the right two plots), and partially shared components associated with technologies (bottom panel: bulk on the left two plots and single cell on the right two plots) by 2s-LCA.

### S.7 Additional Results on Applying SLIDE to Real Data

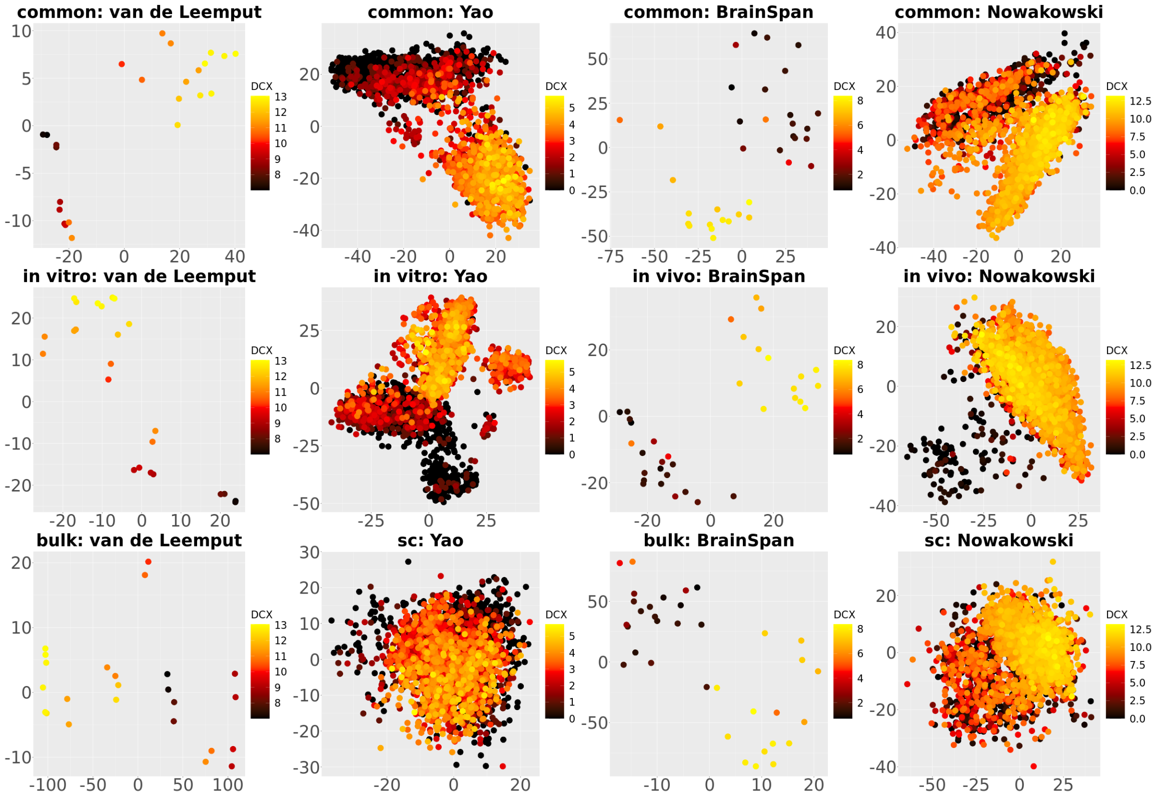


Figure S.27: Scatterplots of scores of each data set corresponding to common components (top panel), partially shared components associated with environments (middle panel: *in vitro* on the left two plots and *in vivo* on the right two plots), and partially shared components associated with technologies (bottom panel: bulk on the left two plots and single cell on the right two plots) by SLIDE.

### S.8 Additional Results for the Validation Study

For those data sets, raw RNA-Seq reads were downloaded from the Sequence Read Archive at the National Center for Biotechnology Information (NCBI). These were analyzed using the Rail-RNA spliced aligner [5] and outputs were fed to the recount2/Snaptron software system [2, 6] to generate the gene-by-sample count tables used here.

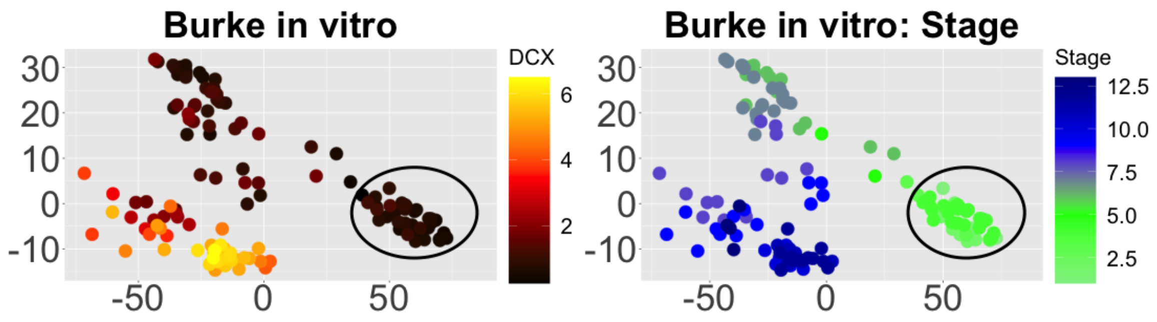


Figure S.28: Projection of an additional *in vitro* bulk RNA-seq data set onto the components shared only by the *in vitro* data sets. Black circles indicate pluripotent stem cells, which are present only in the *in vitro* data sets. Stages: 1-3 (pluripotency), 4-9 (neural progenitors), 10-13 (neurons).

[3] Rick Durrett. *Probability: theory and examples*, volume 49. Cambridge university press, 2019.

[4] Zheng Tracy Ke, Yucong Ma, and Xihong Lin. Estimation of the number of spiked eigenvalues in a covariance matrix by bulk eigenvalue matching analysis. *Journal of the American Statistical Association*, (just-accepted):1–45, 2021.

[5] Abhinav Nellore, Leonardo Collado-Torres, Andrew E Jaffe, José Alquicira-Hernández, Christopher Wilks, Jacob Pritt, James Morton, Jeffrey T Leek, and Ben Langmead. Rail-rna: scalable analysis of rna-seq splicing and coverage. *Bioinformatics*, 33(24):4033–4040, 2017.

[6] Christopher Wilks, Phani Gaddipati, Abhinav Nellore, and Ben Langmead. Snaptron: querying splicing patterns across tens of thousands of rna-seq samples. *Bioinformatics*, 34(1):114–116, 2018.

[7] Yi Yu, Tengyao Wang, and Richard J Samworth. A useful variant of the davis–kahan theorem for statisticians. *Biometrika*, 102(2):315–323, 2015.
